# *Cryptosporidium* sex employs a complex male fertilization apparatus that is required for continued infection

**DOI:** 10.64898/2026.10.13.757955

**Authors:** Aurélia C. Balestra, Maria Merolle, Tim Bergner, Mingyang Tang, Nelida Y. Robles-Ramirez, Anitha Ndayiragije, Bethan A. Wallbank, Abigail M. Daniels, Michael Laue, Christian Klotz, Christopher A Hunter, Boris Striepen

## Abstract

Most apicomplexan parasites replicate asexually to colonize their mammalian hosts, and sexually in the context of transmission. In contrast, for *Cryptosporidium* sex is essential to sustain intestinal infection *in vivo* and therefore is a promising opportunity to prevent and treat the disease. However, the mechanism of sex, which occurs within host enterocytes, is poorly understood, limiting the discovery of targets for such intervention. Here we identify an elaborate fertilization apparatus and numerous of its molecular components in male gametes. We discern two elements, one displayed on the apex, that makes intimate contact with host cells containing female parasites, and a second internal structure, that harbors the membrane fusogen HAP2. Ablation of HAP2 by conditional mutagenesis blocked fertilization and oocyst production, rapidly cured infected mice and left them with enhanced resistance to subsequent challenge. Our studies provide insights into a remarkable fertilization mechanism and highlight specific molecules as targets for translation.

## Introduction

*Cryptosporidium* is a leading cause of diarrheal disease worldwide. This apicomplexan parasite is transmitted by a spore-like stage, the oocyst, which is resistant to water chlorination. Outbreaks due to contamination of drinking or recreational water use are common in the United States and Europe, and they have been a stubbornly regular occurrence despite enhanced surveillance and water management efforts following multiple large incidents^1–4^. Those with primary or acquired cellular immune defects are highly susceptible, and severe cryptosporidiosis is often associated with HIV/AIDS or immunosuppressive treatment in the course of organ transplantation or cancer^5^. The impact of this infection is particularly devastating in young children, where it is dramatically exacerbated by malnutrition^6^. In Africa and Asia alone, 7.5 million children under 2 are estimated to be infected each year, resulting in 200,000 deaths^7,8^. Early childhood cryptosporidiosis often has long-term consequences, including stunted growth and developmental delay^9–11^. Despite this substantial global burden, there is no vaccine, and the only approved drug, nitazoxanide, is ineffective in those populations most vulnerable to the infection^12,13^.

The entire *Cryptosporidium* lifecycle unfolds within the intestinal epithelium of a single human (or animal) host^14^. Infection occurs by ingestion of oocysts, from which sporozoites emerge and proceed to invade intestinal cells. Following invasion, the parasite resides in a distinctive ‘intracellular but extracytoplasmic’ niche that protrudes from the apical surface of the enterocyte into the gut lumen^15^. Following a strictly limited number of asexual replication cycles, *Cryptosporidium* commits to sexual development, leading to the formation of male gamonts and female gametes^16^. Each male gamont gives rise to 16 individual male gametes, which egress from their host cell to fertilize the female gamete. Fusion of male and female gametes and their nuclei in the zygote is immediately followed by meiosis (apicomplexans are haplonts) and sporulation produces new oocysts.

In apicomplexans responsible for malaria, babesiosis, and toxoplasmosis, parasite sex is linked to transmission and occurs in the non-human part of the lifecycle (mosquitoes, ticks and cats, repectively^15,17,18^). However, in *Cryptosporidium,* recent studies have suggested that sex is also required to maintain the infection within the same host^16,19^. The parasite’s lifecycle is hard-wired to transition to sex after only two days^16,19,20^, and continued infection thus depends on a developmental reset provided by sex. This unique biology highlights *Cryptosporidium* sex as a bottleneck and a promising target for therapy and prevention. However, lacking mechanistic understanding of parasite sex currently severely limits the ability to identify and develop targets for specific interference by antibodies or small molecules.

Here, we discovered an elaborate fertilization apparatus localized to the apical end of the male gamete. Structurally, this apparatus is composed of discrete inner and outer elements. We define numerous new molecular components that we suggest act in a two-step process of attachment and fusion. We demonstrate that disrupting fertilization by conditional genetic ablation of the male fusion protein HAP2 cures infected animals of cryptosporidiosis. Such ablation limits infection to two days of asexual development and yields protection to subsequent challenge.

## Results

### Extracellular male gametes fertilize intracellular female gametes

Fertilization relies on the encounter and fusion of gametes, which in apicomplexans is well documented for the malaria parasite *Plasmodium,* where it occurs outside of the red blood cell and in the lumen of the mosquito midgut^21–23^. Only a limited number of electron micrographs of *Cryptosporidium* gamete interaction has been reported to date, but most suggest that fertilization occurs while the female remains within the host cell^24–26^. *Cryptosporidium* fertilization can only be observed *in vivo*, and we sought to visualize this process in *C. parvum,* using highly susceptible interferon *(Ifn)γ^−/−^* mice. Mice were euthanized on day 6 of infection, the peak of burden, the small intestine was resected, and segments of the distal ileum were fixed, dehydrated, and embedded (see Methods for detail). We next used serial block-face scanning electron microscopy, which delivered full volumes of 362 parasites. Searching for parasites harboring diagnostic wall forming bodies (WFB)^24^, we identified 69 female cells and distinguished three types: Type I with small WFB, Type II with numerous larger and more osmiophilic WFB, and developing oocysts, which were more electron dense, and showed an oocyst wall and developing sporozoites (Figure 1B, Video S1 shows full volumes). These types represented 6.1, 9.9 and 3.0%, respectively, of all parasites (Figure 1C).

**Figure 1:**
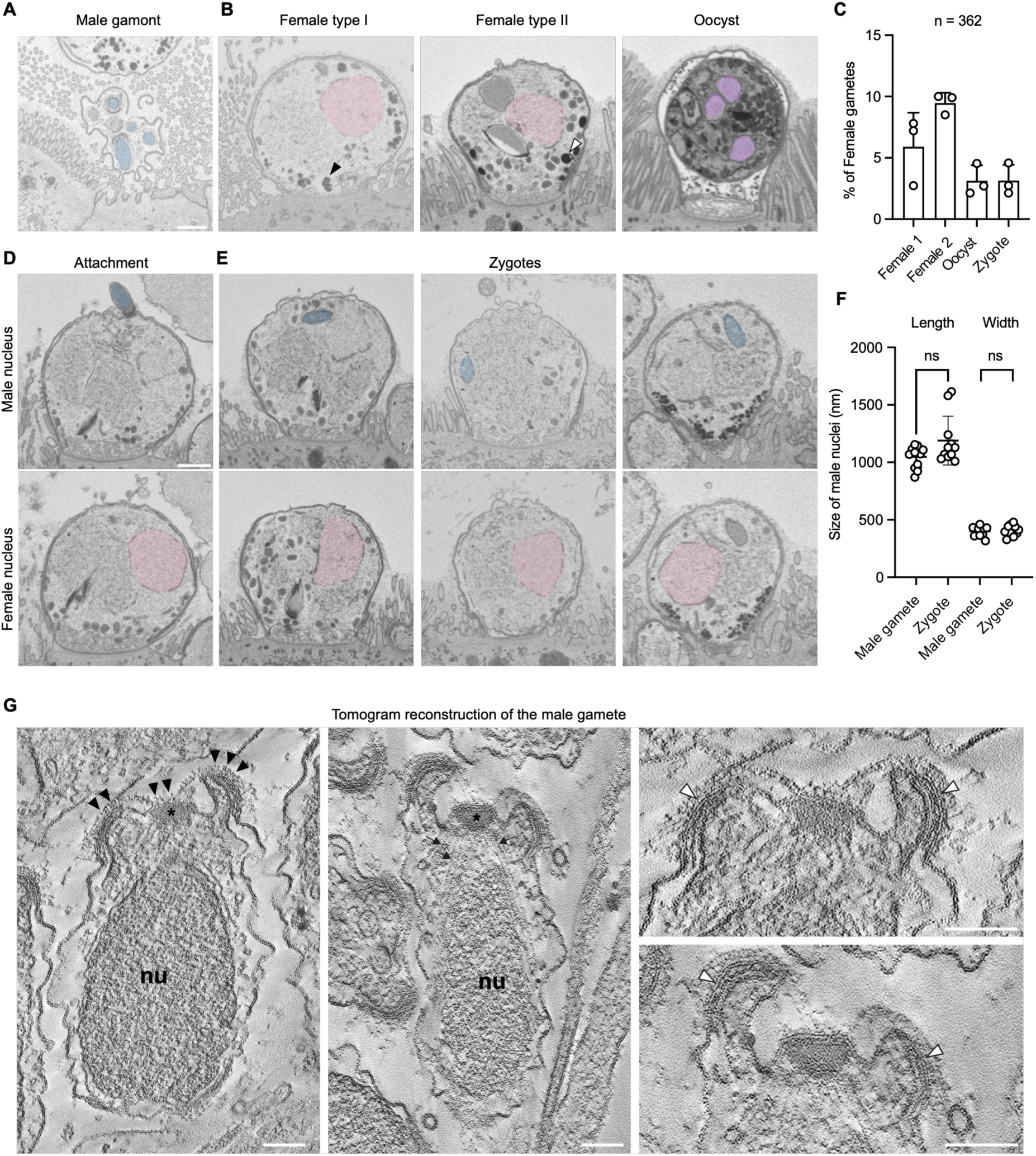
The *Cryptosporidium* male gamete fertilizes the female gamete within its mammalian host cell. (**A, B, D, E**) Serial block-face scanning electron microscopy of intestinal sections from *C. parvum* infected mice. Male nuclei are highlighted in blue, female nuclei in pink, nuclei of developing oocyst in purple. Scale bars = 1 µm. Representative images are shown for male gamonts (**A**) and female gametes at different stages of maturity (**B**). Black arrow: representative WFB for type I, white arrow: larger and more osmiophilic WFB representative from type II. (**C**) Fractions of different stages from the full set of 362 parasite volumes observed. (**D**) A male gamete attached to a host cell containing a female gamete. (**E**) Three zygotes. (**F**) Sizes measure for male nuclei from male gametes or zygotes. Each dot represents a nucleus (n=12 nuclei from male gametes, 11 from zygotes) (Welch’s t test, ns= not significant). (**G**) Electron tomography slices through male gametes within a gamont recorded from intestinal sections. Black arrowheads highlight pronounced densities at the apex, black arrows highlight microtubules and the asterisk indicates a larger electron density which is associated with the apical membrane and tip of microtubules. The white arrowheads indicate the complex multi-layered arrangement of membranes forming the lateral wall of the fertilization pole. Scale bars: 100 nm.

The dataset included 3 male gamonts (a much smaller 0.8% of the population) which contained formed male gametes (Figure 1A, Video S2 shows full volumes). Male gametes are small, polarized cells with a spindle shaped nucleus and an electron dense apical end^24–26^. We observed two female gametes with a male gamete attached to the host membrane covering the parasite (Figure 1D, Video S3 shows full volume). These males were oriented with their apical end towards the female cell. We also identified 11 pronuclear zygotes, intracellular parasites with a large female nucleus and a smaller male nucleus (highlighted in color in Figure 1E, Figure S1, Video S3 shows full volumes of 3 zygotes). Male nuclei were identified by their distinct shape, chromatin texture, and size (Figure 1F). The apical densities noted for free and attached male gametes (Figure 1A and D) were absent from the nuclei observed in zygotes, suggesting that these structures are not internalized during fertilization, or rapidly lost thereafter. Interestingly, all zygotes with an identifiable male nucleus resembled type I female gametes, suggesting a specific point in female development upon which they attain fertilization competence. The more advanced type II stages likely represent zygotes following fusion of the pronuclei.

To visualize the ultrastructure of male gametes in more detail, we used transmission electron microscopy tomography of sections through resin embedded infected tissue (see Methods for detail, Figure 1G). Analysis of tomograms revealed a complex apical pole with a central opening. Longitudinal sections showed an inner chamber that is partially enclosed by arm-like extensions of the cell body which contain multiple membranes (Figure 1G, white arrowheads). This chamber was delimited by a single membrane, with a central protrusion overlaying a large density from which microtubules emanate that enclose the nucleus (Figure 1G, highlighted with an asterisk, microtubules are indicated by black arrows). Notably, the apical extracellular surface of the gamete is covered with a 15 nm thick layer of regularly patterned densities (Figure 1G, black arrowheads). This layer does not cover the central opening in all gametes. We conclude that *Cryptosporidium* fertilization delivers the male nucleus into the intracellular female gamete, and that this interaction involves the apical pole of the male gamete, which has a complex structure.

### Discovering the components of the male fertilization apparatus

To identify components of the fertilization apparatus, we mined the previously established single-cell RNA sequencing dataset^20^. Three expression clusters correspond to male development (Figure 2A), and of the 389 enriched transcripts (p-value <e-50), 58 showed male-unique expression (Figure 2B, Table S1). Transcription factors were enriched in cluster 10, cluster 11 contained the male gamete fusogen HAP2 and the largest share of male exclusive transcripts, and enzymes were enriched in cluster 12 (Figure 2B, Table S1). We prioritized proteins with predicted secretory signal peptides, transmembrane domains, glycosylphosphatidylinositol (GPI) anchors, or domains known to engage in ligand-receptor interactions into an initial list of 10 candidates (Figure S2A). We then used CRISPR/Cas9 guided homologous recombination to either disrupt each of these loci or to introduce epitope tags leaving the genes intact (Figure 2C). *Ifnγ*^−/−^ mice were used to select stable transgenic parasites, oocysts shed with the feces were purified, and genome modification was confirmed by PCR (Figure S2B). While tagged lines could be obtained for all ten genes, knock-out attempts were successful for only two, suggesting that most candidates are likely essential and required for lifecycle completion (Figure S2A).

**Figure 2:**
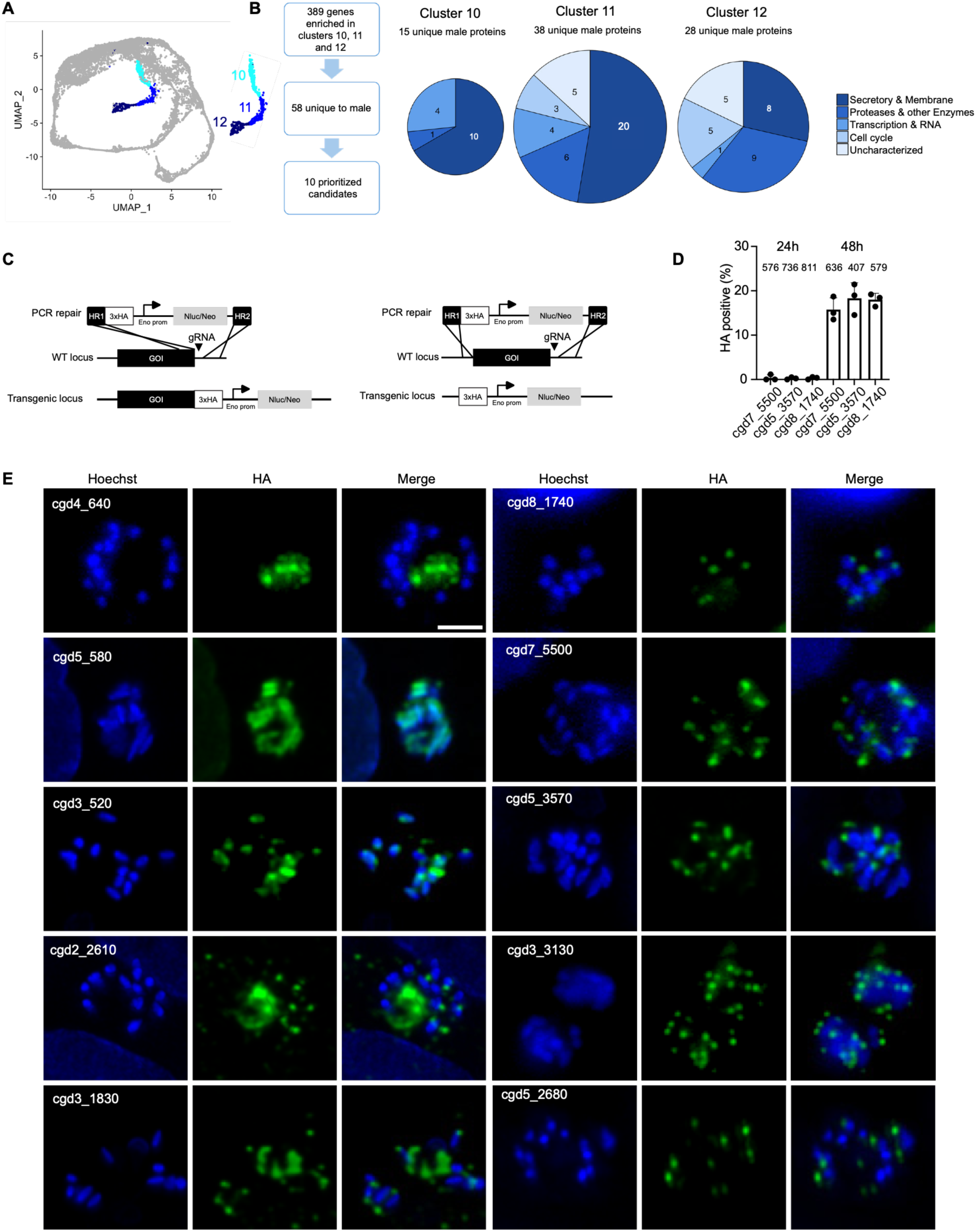
Ten proteins exclusively expressed by male gametes. (**A**) UMAP representation of the *C. parvum* single cell transcriptome^20^ highlighting three clusters representing male development. (**B**) Prioritization scheme for candidate male proteins and annotation of the presumptive biological processes of male exclusive proteins for each of the three distinct expression clusters of male development. Area of each pie chart is reflective of the number of proteins that fall within. (**C**) Schematic maps of gene targeting resulting in HA-epitope- tagged (left) or deleted genes (right). Nanoluciferase (Nluc) and neomycin resistance (Neo) were inserted downstream of the gene.HR: homology region; GOI: gene of interest. (**D**) Quantification by immunofluorescence assay of HA positive parasites at 24h and 48h post-infection in HCT8 cells (3 independent experiments). (**F**) Immunofluorescence micrographs for ten transgenic lines tagging (green) the gene indicated, and showing mature male gamonts (16 bullet-shaped nuclei). Scale bar: 2 µm. Note two proteins localized in the vicinity of the nucleus and seven that label one end of the male gamete.

To investigate protein localization, HCT8 cells were infected with each tagged line followed by immunofluorescence assays at 24h and 48h post-infection to monitor the asexual or sexual phase of parasite development^16,19^. Proteins were engineered to carry a C-terminal 3xHA epitope, except for the GPI anchored cgd3_1830 and cgd3_520, where we used an N-terminal tagging strategy (Figure S2C). The expression of all 10 proteins was restricted to the 48h timepoint and to male parasites (Figure 2E, quantified for three proteins in 2D) and we observed three staining patterns. Protein cgd4_640 was detected in male gamonts, but not in released gametes. Proteins cgd5_580 and cgd3_520 labeled structures proximal to the nucleus of male gametes, and the remaining seven proteins showed polar staining, with two proteins (cgd2_2610 and cgd3_1830) also labeling the residual body of the gamont. Here we focus on the seven proteins with polar localization.

### A family of paralogous proteins forms a multi-membered ring at the apex of the male gamete

*Cryptosporidium* male gametes are very small (0.4 x 1.2 um), and the subcellular features we labeled were close to the diffraction-limit of resolution of the light microscope. To overcome this, we used ultrastructure expansion microscopy (U-ExM)^27^. Briefly, after fixation, cells were embedded in a gel matrix which was allowed to swell isotropically to increase in size by 4.5 times (see Methods section for detail). This technique produced a remarkable improvement and resolved intricate structures. For the first protein analyzed in this fashion, cgd8_1740, we noted a single pronounced ring (green) perpendicular to the bullet shaped male nucleus (blue, Figure 3A). Counterstaining with a N-hydroxysuccinimide (NHS) ester dye to label protein dense structures localized cgd8_1740 to the apical tip of the male gamete (Figure 3E). Applying this technique to additional epitope tagged lines revealed two proteins with similar localization (Figure 3E). All three proteins belong to a family of paralogs named GGC for a shared amino acid motif (see alignment in Figure S3)^20,28^. A fourth member of this family showed similar apical staining by IFA (cgd2_2610, Figure 2E), but labeling was lost in U-ExM limiting the precision of the assignment. We refer to these proteins as GGC1 to 4 (cgd7_5500, cgd2_2610, cgd5_3570 and cgd8_1740, respectively). All of them formed rings, albeit of differing diameter, thickness and height (Figure 3B-D). This may reflect the presence of different individual structures or differentially distributed components of a common feature, with GGC1 present at the periphery. The primary sequence similarity of these four GGC proteins is modest (Figure S3), however modeling using AlphaFold revealed structural conservation with a root mean square deviation of the structural alignment between 1.081 and 1.336 Å (Figure S4A, Video S4). All four proteins are predicted to feature a large, conserved N-terminal domain (Figure 4A), which for GGC1, GGC3 and GGC4 is linked to a smaller more varied domain through a flexible loop. The structure of the larger GGC2 (296 kDA vs ∼165 kDa), revealed what appears to be a duplication of the N-terminal domain in addition to a small C-terminal region again bridged by a flexible linker (Figure 4A).

**Figure 3:**
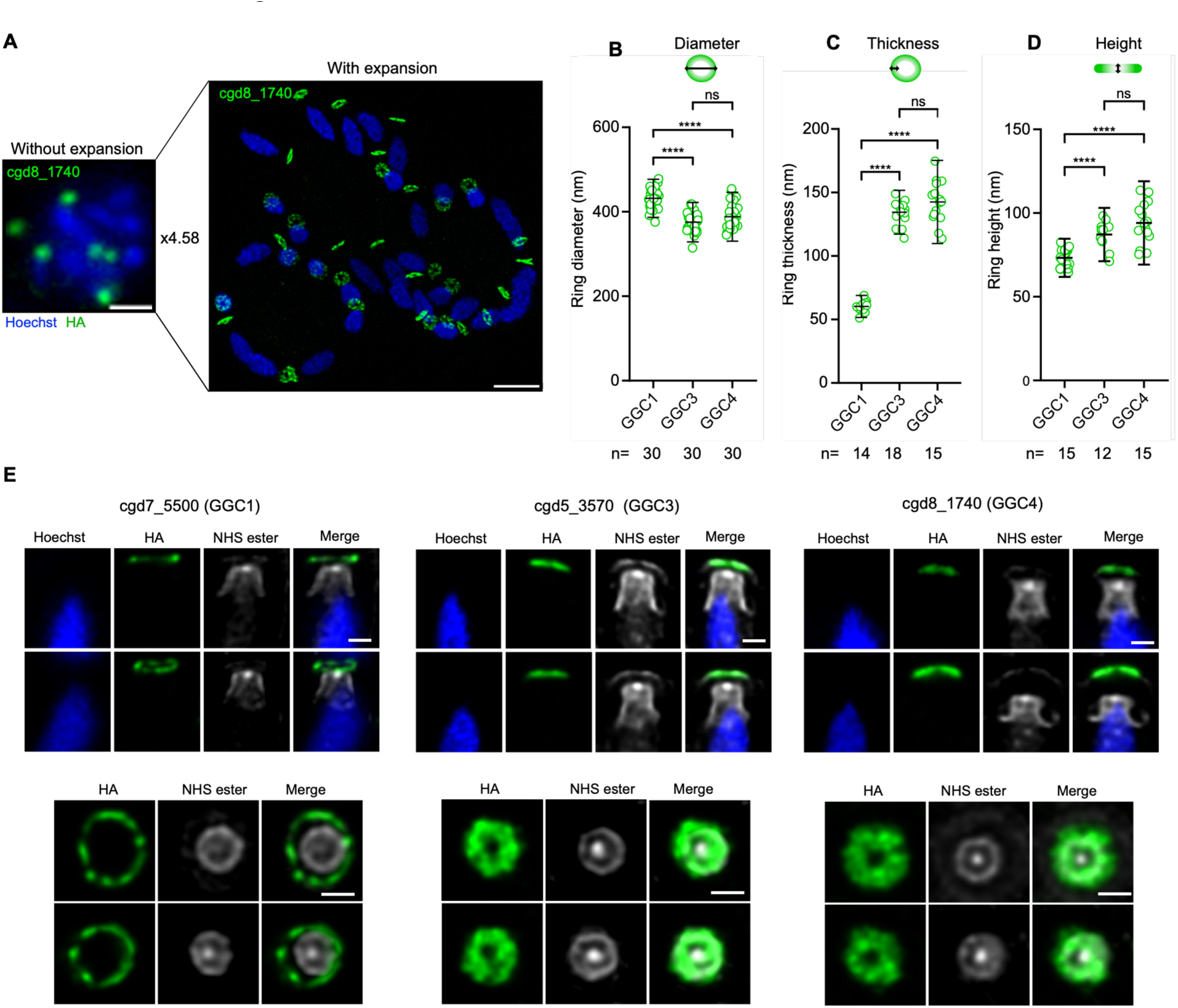
Multiple rings form an outer fertilization apparatus at the apex of the male gamete. (**A**) A representative image showing the localization of cgd8_1740-HA by IFA (left) or by U-ExM (right). Scale bar: IFA - 1 µm; U-ExM - 4.5 µm. Expansion factor: 4.58. (**B-D**) Quantification of the diameter, thickness and height of rings formed by GGC1, GGC3 and GGC4. Each dot represents a male gamete, and the size is divided by the expansion factor of each experiment (ranging from 4.2 to 4.6) (Welch’s t test; ****= p <0.0001, ns= significant). (**E**) U-ExM images of GGC1-HA, GGC3- HA and GGC4-HA. Side (top) and top down (bottom) view of male gametes. HA (green), Hoechst (blue) and N-hydroxysuccinimide ester (NHS) protein counterstain in white. Scale bars: 1 µm. Expansion factor: GGC1: 4.50, GGC3: 4.58, GGC4: 4.58.

**Figure 4:**
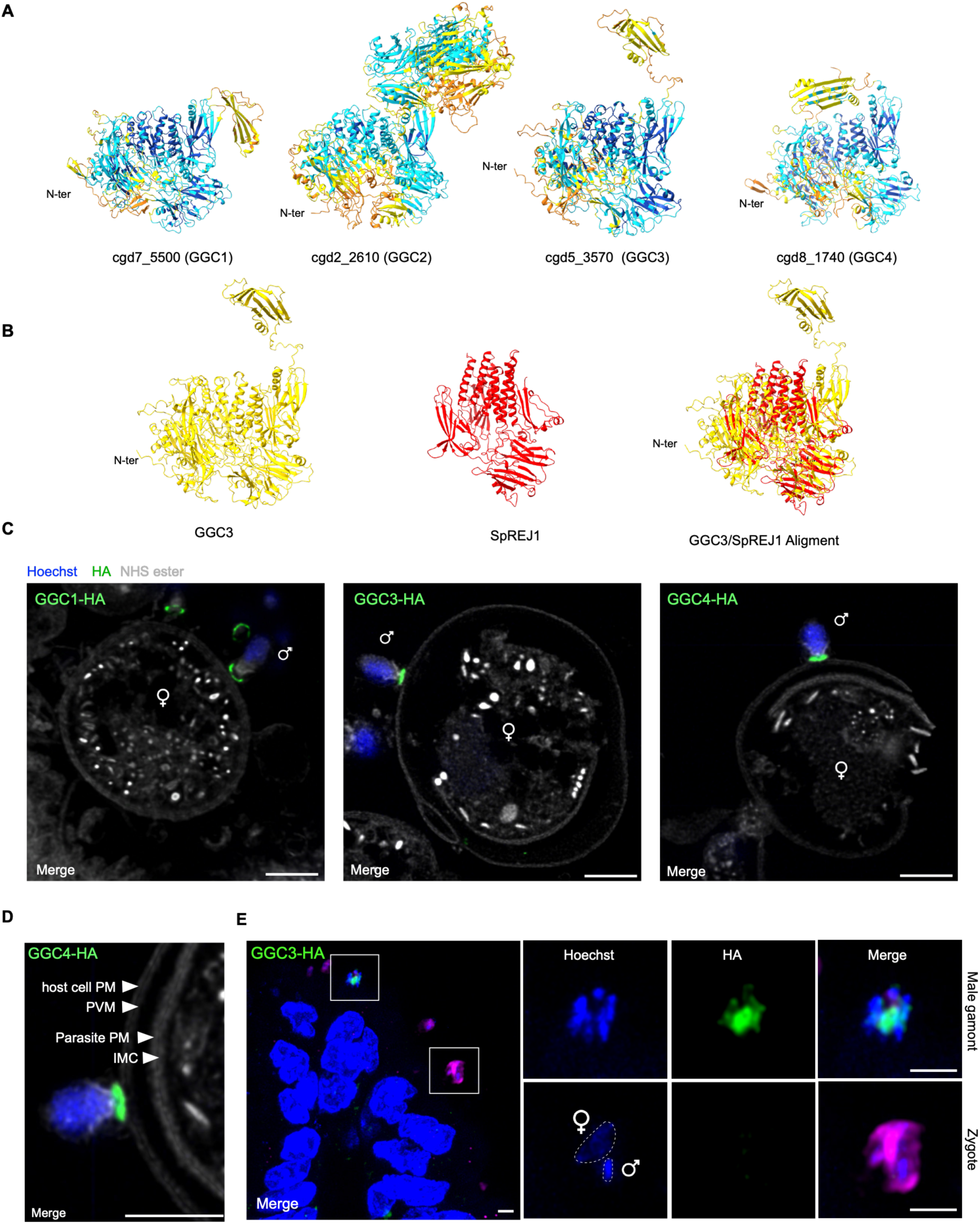
GGC proteins localize at the site of male gamete attachment. (**A**) AlphaFold structural predictions for GGC1, GGC2, GGC3 and GGC4 (omitting the signal peptide). Note conserved N-terminal domain. See Video S4 for movie of overlay. (**B**) Left: AlphaFold prediction of GGC3 and SpREJ1’s REJ domain (SpREJ1_468-1187_). Right: Structure alignment of GGC3 and SpREJ1_468-1187._ RMSD: 1.367 Å across 71 residues. (**C**) U-ExM of infected HCT8 cells showing male gametes attached to the host cell surrounding the female gamete. Scale bars: 4.5 µm. Expansion factor: GGC1-HA: 4.33, GGC3-HA: 4.58, GGC4-HA: 4.50. (**D**) Zoom in of the GGC4-HA from B to highlighting multiple distinct membranes: PM: plasma membrane, PVM: parasitophorous vacuole membrane, IMC: inner membrane complex. (**E**) Representative intestinal sections from GGC3-HA infected mice, with inset showing a male gamont and a zygote. The male and female nuclei of the zygote are highlighted. Magenta: lactate dehydrogenase (LDH) antibody used as pan-parasite stain. Scale bars: 2 µm. Note that GGC3 is readily detected in gamonts but is not present in zygotes.

### GGC proteins localize to the site of gamete interaction

BLAST searches using GGC proteins failed to yield significant orthologs outside of the genus *Cryptosporidium*, however, structure-based searches using FoldSeek revealed similarity to receptor of egg jelly (REJ) domains, originally identified in sea urchin sperm^29,30^ (Figure 4B, Figure S4B-D and Video S4). These domains feature six stereotypically placed cysteine residues, which we note are conserved in *Cryptosporidium* GGC proteins (Figure S4E). REJ domains are found in the ectodomains of membrane proteins and have been linked to receptor function, the sperm acrosome reaction and proteolytic activation^29–33^. Given their structural similarity to sperm receptors, we wondered whether the location of these GGC proteins might facilitate recognition or binding of the female gamete or its host cell. To explore this, we infected HCT-8 cultures with transgenic parasites expressing HA epitope tagged GGC1, GGC3 or GGC4, respectively and fixed cells at 50h. We documented numerous interactions between male and female gametes, and expansion microscopy revealed GGC proteins at the contact site between the gametes (Figure 4C). U-ExM resolved the three membranes surrounding the female gamete with GGC1, GGC3 and GGC4 in tight apposition to the host plasma membrane overlaying the female (Figure 4C-D). We next imaged parasites expressing tagged GGC1 and GGC3 proteins in histological sections from the intestines of infected mice, where fertilization is operational. We readily observe GGC labeling in male gamonts and extracellular male gametes. However, when inspecting zygotes carrying both female and male nuclei, GGC staining was no longer associated with the male pronucleus (Figure 4E and Figure S5). We conclude that the apical ring of the male gamete attaches to the female-containing host cell, making the GGC proteins that assemble into this structure candidates to serve as receptors. Consistent with our earlier electron microscopy studies of zygotes, this apical structure appeared not to be internalized during fertilization.

### The fertilization apparatus includes an internal component

Below the apical ring, U-ExM revealed a previously uncharacterized feature of the male gamete, a protein-dense structure positioned above the nucleus. This structure has a top hat-like cylindrical shape, with a height of 300 ± 60 nm, its diameter is 230 ± 20 nm at the apex to 390 ± 40 nm at the base and is situated 113 ± 35 nm below the apical ring (Figure 5A). NHS labeling also highlights a central focal density at the apex of this structure. This density coincides with the point from which eight perinuclear microtubules emanate, suggesting that it likely contains a microtubule organizing center (MTOC, Figure 5B). We found that three of the male exclusive proteins we epitope tagged appeared as small rings around this central density (Figure 5D-F). Cgd3_3130 contains armadillo repeats (m-ARO, male-armadillo repeats only), and FoldSeek search identified structural similarity with PF16 (Paralyzed Flagella 16, Figure S6A and Video S4). PF16, also known as sperm-associated antigen 6, is an ARO protein conserved across protists, animals, fungi and plants^34^. It is present in male gametes and associated with microtubules in *Plasmodium, Chlamydomonas* and in mice^34–36^. Cgd3_1830 features not only an N-terminal signal peptide but also a C-terminal hydrophobic sequence predicted to act as signal for replacement with a GPI anchor. Cgd5_2680 encodes a protein with five transmembrane domains. Thus, at least two of the three internal proteins are membrane proteins. Considering our electron microscopic insights from Figure 1, we conclude that these proteins localize to the membrane that overlays the central density and forms the recessed portion of the fertilization apparatus. We thus named these proteins Fertilization Apparatus Inner component 1 and 2 (FAi-1 and 2).

**Figure 5:**
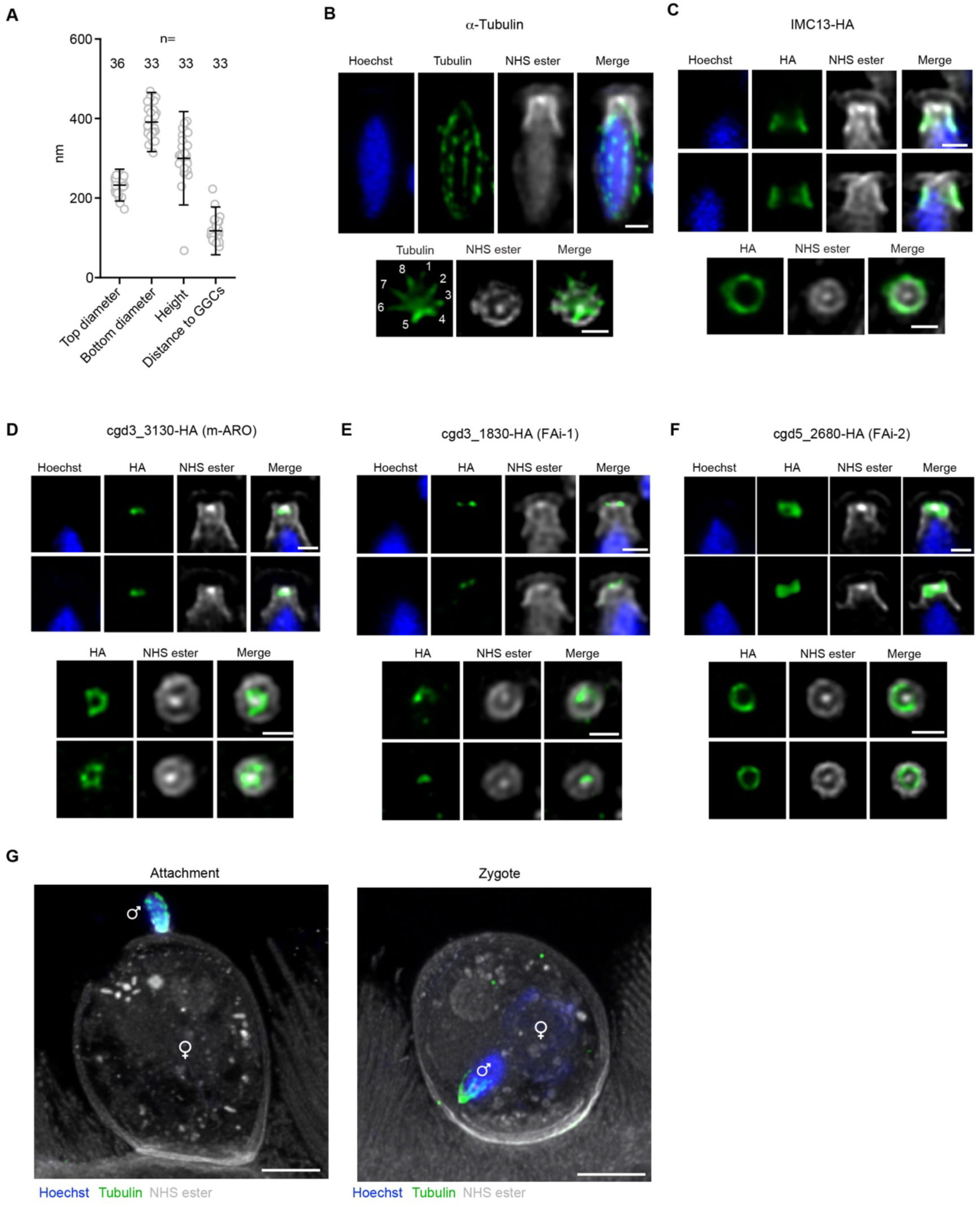
U-ExM reveals an inner component of the fertilization apparatus. (**A**) Quantification of the size of the nuclear hat from 3 independent experiments in HCT8 cells. Each dot represents one male gamete, and the size is divided by the expansion factor of each experiment (ranging from 4.20 to 4.60). (**B**) U-ExM of WT *C. parvum* in HCT8 cells, stained with an α-tubulin antibody. Male gametes shown from the side (top) and top down (bottom). Scale bars: 1 µm. Expansion factor: 4.35. Note the eight microtubules visible in top-down view. (**C**-**F**) U- ExM of IMC13-HA (**C**), m-ARO-HA (**D**), FAi-1-HA (**E**), and FAi-2-HA (**F**) in infected HCT8 cells. Scale bars: 1 µm. Expansion factor: C: 4.25, D: 4.50, E: 4.58, F: 4.33. (**G**) U-ExM of intestinal sections showing a male gamete attached to the host cell surrounding the female gamete (left) and a zygote (right), stained with an α-tubulin antibody. Note that the tubulin is internalized with the male nucleus in the zygote. Scale bars: 5 µm.

We wondered whether the lateral portion of the hat-like structure might be related to the inner membrane complex (IMC). The IMC is a double-membrane cylinder that underlies the plasma membrane of apicomplexans, ciliates and dinoflagellates, which is critical to cell shape and motility^37^. The rigidity of this structure rests on intermediate filament-like proteins called alveolins. Of the 81 *Cryptosporidium* proteins assigned to the IMC/plasma membrane cluster by spatial proteomics^38^, seven are orthologs of *Toxoplasma* alveolin proteins, and the transcripts of five are detected in male gametes of *Cryptosporidium* (Figure S6B). Cgd5_4080, an ortholog of TgIMC13^39^, is highly transcribed in merozoites and male gametes (Figure S6C). We epitope tagged CpIMC13 in *Cryptosporidium* (Figure S6D-E) and found that in merozoites, CpIMC13 mirrors the localization to the basal complex previously found for *Toxoplasma* tachyzoites (Figure S6F). In male gametes, CpIMC13 is indeed localized at the sides of the nuclear hat (Figure 5C). We wondered whether some of these elements may be translocated to the female along with the nucleus during fertilization. We analyzed the localization of IMC13 and male tubulin prior and post fertilization in sections from infected mice. Both are observed in attached male gametes, but only the microtubules are found in zygotes (Figure 5G and Figure S7A-B). Overall, we identify an apparatus with at least two components: a ring system on the apical surface of the gamete made up of multiple GGC proteins, and an internal membranous component harboring distinct membrane proteins surrounding a central MTOC.

### The fusogen HAP2 localizes to the internal component of the fertilization apparatus

Hapless 2 (HAP2) is a membrane protein that closely resembles class II viral fusogens and is critical to fertilization in plants and algae, some animals, and many protists including the malaria parasite *Plasmodium*^40–42^. HAP2 was the first male protein identified in *Cryptosporidium*^19^. Having identified distinct structures at the apical end of the male gamete, we wondered where among those HAP2 would be found. Using a parasite strain in which we tagged HAP2 with a HA epitope, we analyzed male gametes by expansion microscopy. To our surprise, we found HAP2 was not localized at the apex of the gamete but at the center of the internal membrane (Figure 6A), where it formed a ring similar to those observed for FAi-1 and FAi-2. We next imaged HAP2 during male and female interaction in HCT-8 cultures. HAP2 was found at the gamete pole closest to the female, but remained in its recessed position, and was not in direct contact with the membranes of either host cell or female gamete (Figure 6B). As noted, *Cryptosporidium* fertilization does not occur in culture, and we conclude that this block lies downstream of initial recognition and GGC protein interaction, yet upstream of HAP2 mediated fusion. Topologically male and female membranes are distant and separated by male, female and host structures including two membranes. Overall, this suggests a multistep mechanism of fertilization that repositions HAP2 following initial recognition. How the gametes accomplish this repositioning remains unknown.

**Figure 6:**
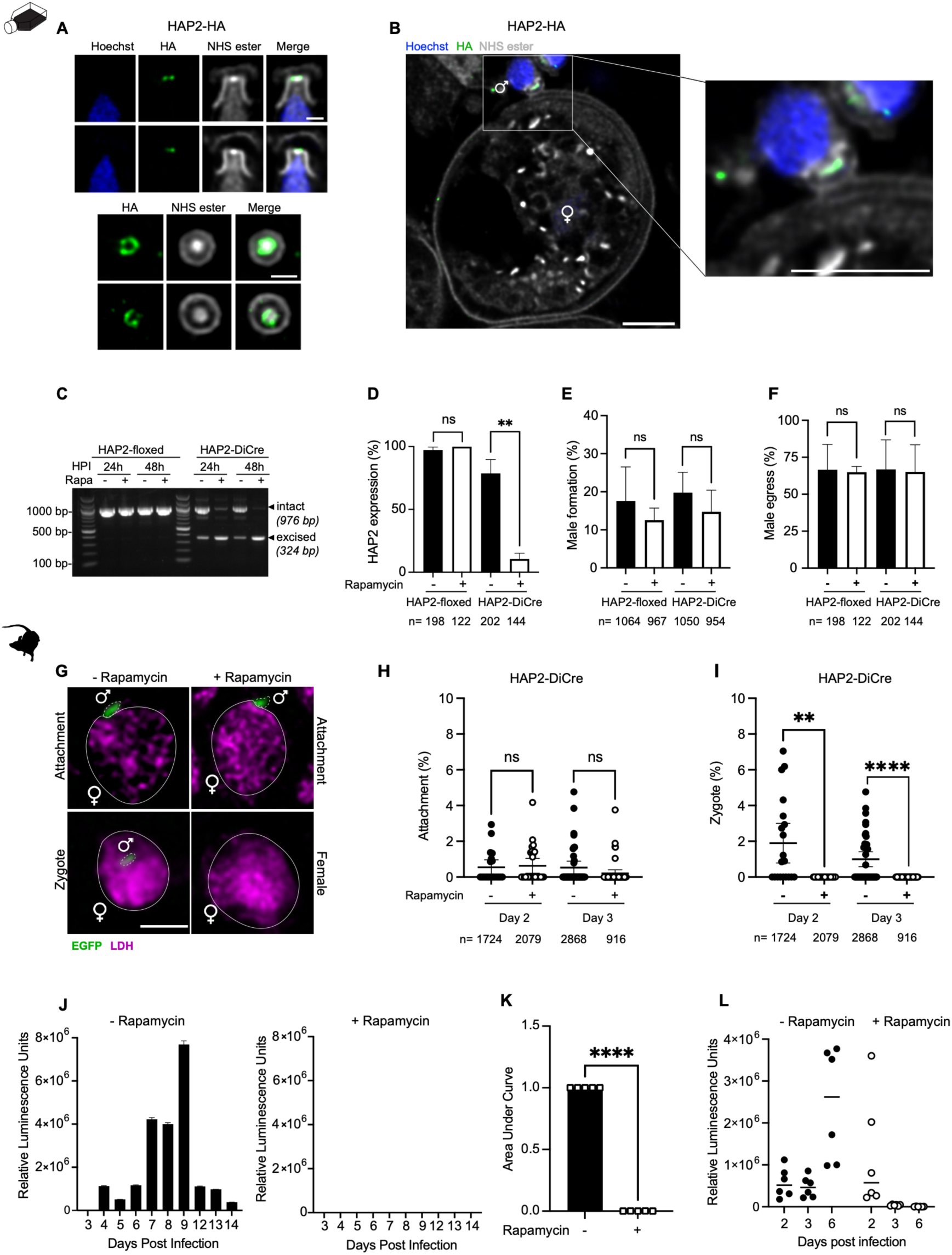
HAP2 is essential for fertilization and required for continued infection. (**A**) U-ExM of HAP2-HA in HCT8 cells. Male gametes shown from the side (top) and top down (bottom). Scale bars: 1 µm. Expansion factor: 4.41. (**B**) U-ExM of HAP2-HA in HCT8 cells showing a male gamete attaching to a host cell harboring a female. Inset shows detail of membranes at the contact. Scale bars: 5 µm. Expansion factor: 4.50. Note that HAP2 consistently does not contact host or female membrane. (**C**) Diagnostic PCR showing inducible excision of 652 bp floxed HAP2 sequence. HCT8 cells infected for 24h or 48h with HAP2-floxed or HAP2-DiCre were grown in presence of vehicle or rapamycin. (**D**-**F**) HCT8 cells were infected with HAP2-floxed or HAP2- DiCre for 48h and treated with rapamycin or vehicle (Welch’s t test; **: p < 0.01, ns: not significant, n= total number of males counted per condition for D and F, total number of parasites for E). (**D**) HAP2 expression in male gametes was assessed by quantifying the number of HA positive 16N parasites. (**E**) Male gametogenesis was assessed by quantifying the number of 16N parasites compared to the total number of parasites (assessed by VVL staining, see representative image in Figure S8E). (**F**) Male egress was assessed by quantification of the number of 16N parasites with at least 1 nucleus outside of the VVL staining (representative image in Figure S8E). (**G**) Representative images of tissue sections from the intestine of mice infected with HAP2-DiCre, treated with vehicle or rapamycin. Top: Male gametes are attached to female gametes. Bottom: in absence of rapamycin, zygotes with internal male nuclei are readily observed. In presence of rapamycin, no zygotes were detected, all female parasites observed were devoid of male nuclei. EGFP: male nuclei; LDH: pan-parasites stain. Scale bar: 2 µm. (**H**-**I**) Quantification of tissue sections as shown in **G**. (Welch’s t test; **: p < 0.01,****: p < 0.0001, ns: not significant, n=total number of parasites scored per condition). Male attachment (**H**) and zygotes (**I**) shown as percentage of all parasites, counted by LDH staining. Each dot represents a field of view from a total of 14 mice. (**J**) Luminescence assay measuring fecal oocyst shedding of infected mice treated with vehicle (left) or rapamycin (right). (**K**) Normalized area under the curve of oocyst shedding (as seen in **J**) of 5 independent experiments. (Welch’s t test; p < 0.0001, n= 30 mice). (**L**) Luminescence assay of intestinal punch biopsies from infected mice treated with vehicle or rapamycin. Note early colonization due to unimpeded asexual growth under rapamycin and lack of reseeding after 3 days. Each dot represents one biopsy, 12 mice total.

### Conditional ablation of HAP2 blocks fertilization

To study the function of HAP2, we generated a conditional knock-out mutant using an inducible cre recombinase (DiCre)^43^. In this system, rapamycin induces reconstitution of active recombinase, leading to the excision of loxP-flanked sequences. We first constructed a “HAP2- floxed” strain in which the last 467 bp of the gene, including those encoding the transmembrane domain, were flanked by loxP sites and we appended a 3xHA tag (Figure S8A). To facilitate downstream phenotypic studies, the locus was also engineered to express a copy of Histone 2B fused to GFP which we placed under the control of a male-exclusive promoter (cgd4_4380) to exclusively mark male gametes with a green nucleus (Figure S8A-B). A second strain expressing DiCre recombinase was equipped with tdTomato under the control of the COWP1 promoter to mark the cytoplasm of female parasites (Figure S8A and C). We then crossed these two lines by co-infecting *ifnγ^−/−^* mice, and recombinant parasites were selected by treatment with paromomycin and BRD7929^28,44^. The presence of both modified loci was validated by PCR mapping (Figure S8D), and we refer to this line as HAP2-DiCre.

We next infected HCT8 cells with the HAP2-floxed and HAP2-DiCre strains and treated with ethanol vehicle or rapamycin for 24 and 48h. Genomic DNA was extracted and the HAP2 locus was amplified by PCR. As expected, the locus remained intact in the floxed parasites lacking the recombinase. In the HAP2-DiCre line, rapamycin treatment resulted in efficient excision, which was near complete after 48h. Some excision occurred in the absence of rapamycin due to leaky DiCre activity as previously reported^43^ (Figure 6C). To investigate the impact on HAP2 protein, cultures were fixed at 48h and subjected to IFA using an HA antibody. In the HAP2-floxed line, 97 ± 1.9.% of males were HA-positive. In the HAP2-DiCre line treated with vehicle, 78.7 ± 9.0% of males showed staining, which dropped to 10.5 ± 3.7% in those treated with rapamycin (Figure 6D).

Next, we investigated the impact of loss of functional HAP2 on the biology of male gametes. All parasites were detected by *Vicia villosa* lectin (VVL) staining, mature male gamonts by their 16 nuclei, and those with nuclei located outside of VVL-stained areas were scored as egressed (representative images are shown in Figure S8E). No significant difference in the number of males formed (Figure 6E) or their ability to egress (Figure 6F) was observed. To understand whether HAP2 deficient gametes were still fertilization competent we turned to *in vivo* experiments. *Ifnγ^−/−^* mice were infected with HAP2-DiCre parasites, treated with vehicle or rapamycin by oral gavage beginning on day five of infection, and killed after two or three days of treatment. The distal ileum was resected, processed for histological analysis (see Methods section for detail), and tissue sections were analyzed by immunofluorescence. Zygotes were identified by the presence of a distinct male and female pronucleus in the same parasite cell as shown in Figure 6G. In vehicle treated mice, we detected 65 zygotes among the 4,592 recorded total parasites (1.4%). In stark contrast, no zygotes were observed among 2,995 parasites in rapamycin-treated mice (Figure 6H- I, 3 independent biological experiments, 14 mice total). Note that attachment of male gametes remained intact, and consistent with our tissue culture experiments, we readily identified male gamonts and gametes in these sections. We conclude that HAP2 is not required for male gametogenesis, gamete egress, or attachment but essential for gamete fusion.

### Blocking fertilization by ablation of HAP2 cures infected mice

Following fertilization, zygotes enter meiosis and sporulation, which results in the production and release of oocysts. *Ifnγ^−/−^* mice were again infected with HAP2-DiCre parasites and treated from the beginning of infection with vehicle or rapamycin by oral gavage for 5 days. Oocyst shedding was monitored daily by measuring fecal luminescence^45^. The feces of vehicle-treated mice showed robust luciferase activity typical for the course of infection in this model, however, rapamycin treatment ablated fecal luciferase activity (Figure 6J). Figure 6K shows normalized area under the curve comparisons for 5 independent biological experiments (30 mice total, p<0.001). To rigorously assess the impact of HAP2 loss not only on oocyst production but also on continued asexual development and intestinal colonization, we measured parasite burden directly in the intestinal epithelium. Mice were infected with HAP2-DiCre, treated daily with vehicle or rapamycin, and sacrificed on days 2, 3 and 6 post-infection. The distal ileum was resected, and luminescence assays were performed on tissue biopsy punches. In vehicle-treated mice, we observed luciferase on all days and an increase in signal over time (Figure 6L, Figure S8F). In rapamycin-treated mice, parasites were readily detectable on day 2, much reduced on day 3 and undetectable on day 6 (Figure 6L, Figure S8F). We conclude that the mutant successfully runs through the typical 2-day asexual portion of the lifecycle (as observed in culture) but then fails to complete the sexual phase to reset the lifecycle for the next round, and the infection dies out. Ablating sex thus cures the infection.

### Infection attenuated by blocking sex induces protective immunity

Our studies demonstrated that genetic ablation of HAP2 allowed brief asexual colonization before ending the infection. We thus asked whether blocking parasite sex impacts the susceptibility of mice to subsequent infection. To establish a suitable dose, *ifnɣ^-/-^* mice were infected with either 5,000 or 50,000 HAP2-DiCre oocysts and rapamycin was orally administered daily for five days. This regimen produced a low burden infection that was cleared by day 5 and produced no clinical signs of disease (Figure 7A). Mice were challenged with wildtype *C. parvum* parasites carrying an Nluc marker 30 days after confirmed resolution of the primary infection. Uninfected mice that had received the same initial rapamycin treatment served as naïve controls. Upon challenge, mice that had initially received 5,000 HAP2-DICre oocysts exhibited parasite burdens comparable to naïve controls (Figure 7B). Those that had received 50,000 oocysts showed diminished secondary infection and exhibited faster control with a burden reduced by two orders of magnitude at the peak of infection (Figure 7B). We therefor used 50 000 oocysts in subsequent experiments. Across 6 independent vaccination experiments, area under the curve analysis showed a consistent and significant decrease in parasite burden when comparing to naïve mice (Figure 7C). In two of these experiments, mice were rechallenged with a highly virulent mouse-adapted strain of *C. parvum*^46^ (shown in purple in Figure 7C). Histological studies were conducted on the small intestines of mice eight days following rechallenge with mouse-adapted *C. parvum.* Naïve mice presented with a highly parasitized epithelium and the signs typically associated with cryptosporidiosis including villus blunting and crypt hyperplasia (Figure 7E-F). In contrast, vaccinated mice showed few signs of infection, with normal crypt depths (Figure 7G) and much reduced parasite numbers per villus (Figure 7H). *Cryptosporidium* causes severe chronic disease in individuals with deficient cellular immunity including those suffering from HIV/AIDS^5^, we thus wondered whether CD4^+^ T cells were also important to the protection observed here. We depleted CD4^+^ T cells immediately prior to rechallenge by antibody treatment and then compared parasite burdens to naïve mice, or those that were vaccinated and subjected to T cell depletion. While vaccinated mice were again protected, the CD4-depleted group was highly susceptible to rechallenge, with high parasite burdens and all mice succumbing to infection (Figure 7D). We conclude that limiting the infection by preventing parasite fertilization provides non-sterile protection to mice reducing parasite burden by a T cell-dependent mechanism.

**Figure 7:**
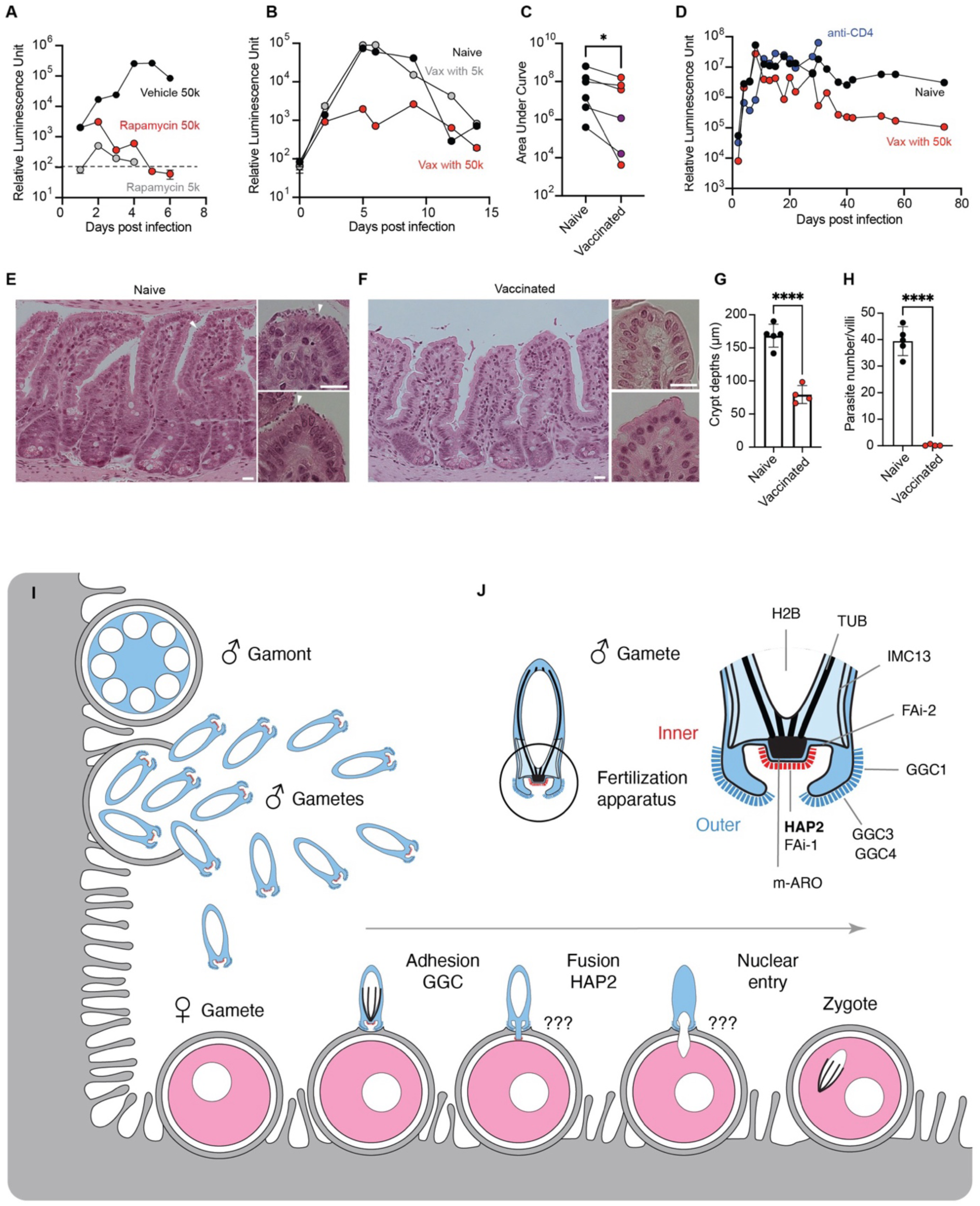
Infection attenuated by loss of fertilization induces protective immunity. (**A**) Luminescence assay of mouse feces showing oocyst shedding of primary infection. Mice were infected with 50,000 HAP2-DiCre oocysts and treated with vehicle or rapamycin or infected with 5,000 HAP2-DiCre oocysts and treated with rapamycin. (**B**) Fecal luminescence assay following rechallenge with 10,000 oocysts of a *C. parvum* Nluc/neo strain of either naïve mice (not infected but treated with rapamycin) or mice following primary infection with 5,000 and 50,000 HAP2- DiCre oocysts. (**C**) Area under the curve of 6 independent experiments as shown in B - Vax with 50,000 oocysts. Mice were rechallenged with 10 000 oocysts of a *C. parvum* Nluc/neo strain (red), or mouse adapted *C. parvum* strain (purple) (Wilcoxon signed-rank test, p < 0.05). (**D**) Fecal luminescence assay of *C. parvum* Nluc/neo rechallenge of naïve mice, vaccinated mice, or vaccinated mice treated with anti-CD4. (**E-G**) Mice were vaccinated and rechallenged with 50,000 oocysts of a mouse adapted *C. parvum* strain. Histology of the ileum of naïve (**E**) or vaccinated (**F**) mice on day 8 of rechallenge with mouse adapted *C. parvum*. White arrows highlight parasites. (**G**) Number of parasites counted per villus. Each dot corresponds to a mouse and represents counts from 3 villi for each mouse (Welch’s t test, p < 0.0001). (**H**) Quantification of the crypt depths. Each dot corresponds to a mouse, the crypt length of 15 villi is averaged for each mouse (Welch’s t test, p < 0.0001). (**I**) Working model of *Cryptosporidium* fertilization. Male gamonts (blue) develop and release male gametes into the gut lumen. Male gametes attach to host cells containing female gametes (pink) using the exterior component of the fertilization apparatus. A conformational change enables HAP2 dependent fusion with the female, then nucleus and microtubules translocate, yielding a zygote. (**J**) Model of the male fertilization apparatus highlighting the localization of the proteins identified in this study.

## Discussion

Eukaryotes have evolved fascinatingly diverse ways to reproduce sexually^47^. The high cost of sex and its associated complex mechanisms is balanced by the gain of genetic diversity among recombinant offspring, and the opportunity to repair damaged DNA during meiosis^48^. The importance of these benefits differs among eukaryotic taxa, resulting in varied emphasis on sexual reproduction. Sex can be obligate to reproduction, facultative, switching between asexual and sexual cycles, occur only in response to environmental challenge, or be cryptic yielding almost clonal populations. In apicomplexan parasites reproduction is primarily asexual, and sex is typically associated with transmission. The discovery that *Cryptosporidium* must undergo sexual replication every two days was therefore surprising, and significant, as it revealed a potential vulnerability. Blocking sex by parasite contraception may offer ways to treat or prevent an infection for which we still lack effective drugs and vaccines^8^. Previous studies highlighted the importances of male and female gamete transcription^20,49^ and mRNA homeostasis^50^ for the production of infectious oocysts. Here, we provide the most direct evidence yet, linking the success of parasite fertilization not only to oocyst shedding, but also directly to colonization of the intestinal epithelium (Figure 6J-L and Figure S8F). Such a block limits the infection to two days of initial asexual expansion. Interestingly, this attenuated infection is sufficient to induce non- sterile disease protection. Vaccines using genetically attenuated parasites have been shown to be protective against malaria in mice^51,52^ and humans^53,54^. Similarly, anti-malarial treatment that allowed for a limited window of parasite replication yielded immune protection – an approach dubbed chemovaccination – that has been tested in animals and humans^55–58^. *Cryptosporidium* gametes express stage-specific enzymes, receptors, and ion channels belonging to classes previously shown to be druggable with small molecules. One example is the male exclusive cyclic nucleotide regulatory system believed to control fertilization^14,20^. The pharmacology of phosphodiesterases is well developed and has yielded numerous medicines^59^, and structural differences between apicomplexan and human phosphodiesterases have already been exploited^60,61^ including in *Cryptosporidium*^62^.

Antibodies offer another well-established route to block fertilization, and contraception by vaccination has been studied extensively in mammals^63,64^. Fertilization relies on the interaction of ligands and receptors presented by male and female gametes, and both can be targeted. Immunizing female mice against the sperm receptor Izumo1 led to a marked decrease in fertility^65–68^. Antibodies to its female counterpart Juno similarly block fertilization^69^. Furthermore, immunizing guinea pigs with PH-20, a sperm protein that digests hyaluronic acid present on the outside of the egg^70^, led to infertility in both males and females^71^. In Apicomplexa, targeting sexual stages with the goal of transmission blocking malaria vaccines has been explored widely^72^. Immunization against the gamete receptor Pfs230 has shown particular promise, and a Pfs230 based vaccine has progressed to clinical trial in Mali^73,74^. Importantly, targeting HAP2 with antibodies similarly reduced *Plasmodium* transmission^75–77^.

While *Cryptosporidium* sex is thus of great translational significance, we lack the mechanistic understanding of fertilization to direct our efforts to block it. This study revealed a complex fertilization apparatus at the apical pole of the male gamete in molecular and cell biological detail. Identifying and localizing numerous male proteins we distinguish two components: an outer assembly of rings that enable adhesion, and an internal component that mediates fusion. We demonstrate that HAP2 is essential to fusion and fertilization in *Cryptosporidium.* Without it, male gametes form, egress and attach to cells harboring female gametes, however fertilization fails and no zygotes are detected. These results are consistent with HAP2’s central role in fertilization, which has been most intensively studied in plant and algal model systems^78^. Crystal structures of HAP2 have been resolved for *Chlamydomonas* and *Arabidopsis*^40,79,80^, revealing their similarity to viral class II fusion proteins, which fold into articulated rods composed of three β-sheet-rich domains^40,81^, and trimerization is a prerequisite of activity^40,82^. Viral class II fusogens fold as heterodimers with a partner protein which ensures their proper trafficking and localization^83–88^. Importantly, the partner also acts as an inhibitor, preventing premature fusion by concealing the fusion loop^87,89,90^. Such partners have also been identified for HAP2 in *Arabidopsis*^91^ and *Chlamydomonas*^92^, and it has been suggested that interaction with specific female ligands releases HAP2 from inhibition to trigger gamete fusion. While such inhibitors have not yet been identified in *Cryptosporidium*, their existence is likely, and the gamete transcriptome highlights potential candidates^20^ and we already identified multiple proteins that share HAP2’s localization (Figure 5D-F).

To our initial surprise, we found HAP2 to localize to the internal component of the fertilization apparatus, which does not make immediate membrane contact upon attachment (Figure 6B). This suggests that the male gamete must undergo a conformational change to enable HAP2 contact and fusion. Such changes in the subcellular morphology of male gametes are well documented and often triggered by contact with female gametes^78,93,94^. For example, in *Chlamydomonas,* ciliary attachment activates a signaling cascade that leads to the outgrowth of an actin-filled mating tubule that carries HAP2 at its tip^95–97^. Similarly, the acrosome reaction of animal sperm is an exocytosis event that restructures the sperm head and exposes and relocates critical receptors^98^. In sea urchins, the acrosome reaction is triggered by the interaction of a protein present at the sperm apex, SpREJ1, with a glycoconjugate component of egg jelly^30,31^. This induces actin polymerization at the sperm apex, and extension of a finger-like acrosomal process^93^. In mice, sperm cells lacking PKDREJ, a member of the same protein family^29^, exhibit a delayed acrosome reaction^32^. Strikingly, *Cryptosporidium* GGC proteins, which like SpREJ1 and PKDREJ localize to the apex of the gamete^99,100^ also share structural similarity, the presence of a REJ-like domain. While gamete receptors are often considered fast evolving proteins limiting primary sequence conservation, there are multiple examples of structural conservation. Examples include sperm proteins conserved among vertebrates that bind highly divergent egg receptors^101^, and the shared folding of mollusk sperm receptors and yeast mating proteins^102^.

Figure 7I-J summarizes our current working model of *Cryptosporidium* fertilization. The proteins identified and localized in this study highlight the complex structure of the male gamete. While we find support for roles in adhesion and fusion for GGC proteins and HAP2, respectively, what unfolds in between those events remains to be discovered. How, following initial attachment, does HAP2 reach the membrane of the female gamete and how does it overcome the membranes of the host cell and the parasitophorous vacuole in the process? Interestingly, this is the very step that does not occur in culture, which may hint at a specific contribution of *in vivo* host biology to license or active the male^16,19,103,104^. We hypothesize the extension of a fertilization bridge; however, such a connection has not been directly observed at this point. Finding ways to arrest fertilization either pharmacologically or genetically may make intermediates observable and help to fill this important knowledge gap. While there is evidence that only the male nucleus and its microtubular cage are translocated to the female cell, how that occurs is unknown. Multiple events of molecular recognition and signaling are required to orchestrate the complex sequence shown. These must involve numerous male and female and potentially also host proteins. The single cell transcriptome identified roughly 60 male and 180 female exclusive proteins suggesting a complex but ultimately understandable mechanism. This initial analysis already identified multiple proteins as validated targets for future translation, and continued work is likely to expand that list significantly.

## Material and methods

### Mice

*Ifnγ^−/−^* mice (stock no. 002287; RRID:IMSR_JAX:002287) were obtained from The Jackson Laboratory and maintained as a breeding colony at the University of Pennsylvania. Experimental cohorts consisted of mice aged 4–12 weeks, with age and sex strictly matched across comparison groups. All animal procedures were reviewed and approved by the University of Pennsylvania Institutional Animal Care and Use Committee (IACUC protocol #806292).

### Plasmid construction

All primers used in this study can be found in Table S2. To generate the HAP2-floxed plasmid (pAB003), a synthetic gene fragment was generated by GenScript containing i) a loxP-intron: a loxP site introduced in the cgd1_1320 intron, as described before^43^, ii) a recodonized HAP2 gene fragment, from base pair 2473 to the end of the gene, excluding introns and stop codon (see recodonized sequence in Table S2), iii) a 3xHA tag and the actin 3’ UTR, and iv) another loxP site. This synthetic fragment was introduced into the pLIC-3xHA-Nluc-Neo plasmid using the enzymes AvrII and PstI. To generate the N-terminal floxing and tagging construct (pAB015), the 5’UTR and signal peptide of cgd8_1740 were amplified by PCR. A 5’ loxP and a 3’loxP-intron were added by a second PCR and the construct was introduced into the pLIC-3xHA by Gibson. To generate pCas9- guide plasmids, a guide RNA targeting the gene of interest was annealed with the into the BbsI-digested pCas9 plasmid.

### Maintenance of Cell Lines and Parasite Strains for Infection

Human colorectal adenocarcinoma HCT-8 cells (ATCC CCL-224™) were maintained in RPMI medium supplemented with 10% fetal bovine serum (FBS), 1× penicillin-streptomycin and 1× gentamicin at 37°C and 5% CO₂. Prior to infection, cells were switched to complete RPMI-1640 containing 1% FBS. *C. parvum* oocysts (Iowa II strain, IIa subtype) were obtained from Bunchgrass Farms (Deary, ID). For *in vitro* infection, oocysts were bleached in a 1:4 (v/v) bleach:PBS solution for 3 min at 4°C, pelleted at 16,000 × *g*, and washed once in PBS. Excystation was induced by incubation in 0.2 mM sodium taurodeoxycholate for 15 min at 15°C. Following excystation, sporozoites were pelleted at 16,000 × *g*, resuspended in infection medium, and added to HCT-8 cells.

### Generation of transgenic parasite

Transgenic parasites were generated as previously described^45^. Guide RNAs (gRNAs) were designed using the VEuPathDB CRISPR guide design tool. For C-terminal tagging, a gRNA targeting the 3’UTR of the gene of interest was selected, and the repair template was PCR-amplified from pLIC-3xHA-Nluc-Neo using gene-specific primers with 50 bp of homology arms to facilitate integration. For N-terminal tagging, a gRNA targeting the 5’UTR was chosen, and the repair template was PCR-amplified from pAB015. For floxing of the *hap2* gene, a gRNA targeting the 3’ end of the gene was chosen, and the repair template was PCR-amplified from pAB003.

*C. parvum* oocysts (2.5 × 10^7^) were bleached and washed prior to excystation. Excystation was induced by incubation in 0.2 mM sodium taurodeoxycholate for 10 min at 15°C, followed by a 1- h incubation at 37°C to release sporozoites. Purified sporozoites were resuspended in SF transfection buffer containing 50 μg each of pCas9-guide plasmid and PCR repair template, then nucleofected using a Lonza 4D Nucleofector (program EH100). Following nucleofection, sporozoites were resuspended in cold PBS. Prior to infection, *ifnγ^−/−^* mice received antibiotic treatment for 5–7 days and were administered with 100 μL of 8% sodium bicarbonate via oral gavage immediately prior to infection to neutralize gastric acidity. Mice were infected with 100 μL of transfected sporozoites via oral gavage. Stable transformants were selected by administering 16 mg/mL paromomycin in drinking water. Parasite shedding was monitored by quantifying fecal nanoluciferase activity. Recovered oocysts were purified from feces using sucrose flotation followed by cesium chloride density gradient centrifugation, and stored long-term in PBS at 4°C. Genomic DNA was extracted from purified oocysts using the DNeasy Blood and Tissue Kit (Qiagen) and 5’ and 3’ integration of the cassette was confirmed by PCR for each transgenic strain.

### Parasite cross

To generate the HAP2-DiCre strain, strains were crossed as previously described^44^. Briefly, *ifnγ^−/−^* mice were co-infected with 6000 oocysts of the neomycin-resistant HAP2-floxed strain and 6000 oocysts of the pheRS^r^-Nluc-DiCre BRD7929 resistant strain^44^. Four days post-infection, mice were treated with 16 mg/ml paromomycin in the drinking water and with 10 mg/kg/day BRD7929 by oral gavage for 7 days.

### Rapamycin treatment

Rapamycin (Thermo Fisher Scientific, Waltham, MA) was prepared as a 50 mg/mL stock solution in 95% ethanol. For *in vitro* experiments, rapamycin was added to cultures at a final concentration of 100 nM at the time of infection, with an equivalent volume of ethanol added to vehicle control wells. Genomic DNA was extracted from infected cultures using the DNeasy Blood and Tissue Kit (Qiagen) and used as template for PCR analysis of Cre-mediated excision. For *in vivo* studies, rapamycin stock was diluted in drinking water to deliver 10 mg/kg body weight daily via oral gavage in a volume of 100 μL.

### Measuring parasite burden from feces and tissue biopsies by luciferase assay

To measure oocyst shedding in feces, 20 mg of fecal material was suspended in 1 mL of fecal lysis buffer and homogenized by vortexing for 1 min with 5 glass beads. The supernatant was mixed 1:1 with NanoGlo luciferase assay buffer containing a 1:50 dilution of NanoGlo substrate (NanoGlo Luciferase Assay Kit, Promega, #N1110) in white-bottom 96-well plates, and luminescence was measured using a Promega GloMax plate reader. For intestinal tissue analysis, biopsy samples were obtained from the distal ileum of euthanized mice using a 5-mm biopsy punch. Tissue samples were incubated in 1 mL Nluc lysis buffer for 10 min at 2000 rpm. Luciferase activity in the resulting lysates was quantified using the same protocol as for fecal samples.

### Transmission electron microscopy and electron tomography

To prepare intestinal sections for EM analysis, a fixative solution was made with 4% paraformaldehyde, 2.5% glutaraldehyde in 0.05 M HEPES buffer and the pH was adjusted to 7.2 with NaOH. Mice were killed and the ileum removed, flushed with the fixative solution and cut transversely (pieces <3x3x3 mm). The samples were kept in the fixative for at least 24h. Post- fixation and block staining were carried out sequentially using 1% osmium tetroxide, 0.1% tannic acid, and 2% uranyl acetate. Samples were dehydrated through an ascending ethanol series, infiltrated with ascending epoxy resin/acetone mixtures, and finally embedded in epoxy resin. After sectioning (thickness approx. 150 nm), samples were examined on a transmission electron microscope (Tecnai Spirit, Thermo Fisher Scientific) equipped with a LaB6 filament, operated at 120 kV. Tilt series were recorded using a bottom-mounted Eagle camera (Thermo Fisher Scientific) and the TIA software with the integrated tomography module (Tecnai Imaging and Analysis, Thermo Fisher Scientific). Tomograms were reconstructed with the IMOD software package version 4.7^105^ according a previously described procedure^106^.

### Serial block-face scanning electron microscopy (SBF-SEM)

SBF-SEM preparation of fixed mouse intestinal tissue was performed using an enhanced contrasting protocol including ferrocyanide-reduced osmium tetroxide post-fixation, thiocarbohydrazide-osmium liganding (ROTO), and en bloc staining with uranyl acetate and lead aspartate^107^. Subsequently, samples were dehydrated through an ascending ethanol series, infiltrated with ascending epoxy resin/acetone mixtures, and finally embedded in epoxy resin. SBF-SEM imaging and sectioning were carried out using a field-emission scanning electron microscope equipped with an integrated ultramicrotome (Teneo Volumescope, Thermo Fisher Scientific). The microscope was operated at 2 kV and 100 pA in low vacuum (0.4 mbar), and images were acquired with the Volumescope Directional Backscatter (VS-DBS) detector at a pixel size of 4 nm (frame size 10,240 × 10,240 pixels), a dwell time of 1 µs per pixel, and a section thickness of 40 nm. Image stacks were processed (slice registration, AI-based denoising^108^, anisotropic diffusion filtering, and median filtering) using either Fiji^109^ or Dragonfly 3D World (v2025.1, Comet Technologies Canada Inc.).

### Immunofluorescence Assay on HCT8 cells

HCT-8 cells cultured on 12 mm round coverslips in 24-well plates were infected with excysted oocysts and incubated for 24 h to assess meront development, or 48 h to observe male and female gametes. Infected HCT-8s were washed once in PBS before fixing in 4% paraformaldehyde for 15 minutes. Cells were washed once in PBS before permeabilization with 0.25% Triton X-100 for 15 minutes. Cells were washed once before blocking with 4% bovine serum albumin (BSA) for 30 to 60 minutes at room temperature. Primary antibodies were then added for 1 hour, and for this study included rat anti-HA (1:1000, Roche clone 3F10, #11867423001), VVL-FITC (1:1000, Vector FL1231) and mouse anti α-tubulin (1:1000, Sigma-Aldrich #T6199). Coverslips were washed three times (5 min incubation each time) in PBS and incubated with secondary Alexa Fluor antibodies for 1 hour. Coverslips were then washed once in PBS, then once in PBS containing 1:10,000 Hoechst for 5 minutes, and then once more in PBS before mounting on slides with Fluorogel (Electron Microscopy Science, Hatfield, PA) or Vectashield (Vector Laboratories).

### U-ExM on HCT8 cells

U-ExM was performed based on^28,38,110^. A total of 400,000 excysted oocysts were used to infect HCT-8 cells grown on P24 coverslips. At 24–48 h post-infection, coverslips were fixed in 4% paraformaldehyde (Electron Microscopy Sciences) and incubated overnight at 37°C in a solution of 1.4% formaldehyde (Sigma-Aldrich) and 2% acrylamide (Sigma-Aldrich). Gelation was initiated by adding a polymerization mix composed of 90% monomer solution (18% sodium acrylate, 10% acrylamide, 0.1% *N,N*ʹ-methylenebisacrylamide in PBS), 5% tetramethylethylenediamine (TEMED, 10% stock), and 5% ammonium persulfate (APS, 10% stock) to the coverslips. Gels were polymerized on ice for 5 min, followed by 1 h at 37°C in a humidified chamber. Gels were released from the coverslips and denatured at 95°C for 90 min in denaturation buffer (200 mM SDS, 200 mM NaCl, 50 mM Tris, pH 9). Expansion was achieved by incubation in ddH₂O, followed by controlled shrinking in PBS, blocking in 3% BSA, and overnight incubation with primary antibody. Primary antibodies include rat anti-HA (1:500, Roche clone 3F10, #11867423001), mouse anti-α- tubulin (1:1000, Sigma-Aldrich #T6199). Gels were incubated with secondary antibody for 2.5 h (Hoechst, 1:1000, Thermo Fisher Scientific #H3570; anti-rat 488, 1:500, Thermo Fisher Scientific # A-11006; Atto 647N NHS ester, stock in DMSO at 2 mg/mL, 1:200, Sigma-Aldrich #18373), then re-expanded, and mounted onto poly-D-lysine-coated MatTek 35-mm glass-bottom dishes for imaging.

### Histology

Mice were infected with 10,000 oocysts/mouse unless stated otherwise. Fecal NLuc was monitored and mice were killed when the luminescence exceeded 200 000 relative light units, unless stated otherwise (Day 5-10 post-infection). To prepare intestinal sections, mice were euthanized and the ileum removed, flushed with PBS, cut open longitudinally and ‘Swiss-rolled’, as previously described^111^. Samples were fixed overnight at 4°C in 10% neutral buffered formalin (Thermo Fisher Scientific) followed by 30% sucrose overnight (Thermo Fisher Scientific #15-188- 51). Tissue was frozen in OCT solution (Sakura Finetek), cryo-sectioned (5 to 20 µm sections) and mounted on slides. For histopathologic analysis by hematoxylin and eosin staining, a Gemini Autostainer (Thermo Fisher Scientific) was used to deparaffinize the sections in xylene before rehydrating them by sequential baths in 100% ethanol, 95% ethanol and 70% ethanol followed by running water for 2 minutes. Hematoxylin and eosin stains were applied as previously published^112^. Slides were imaged at 40X and 63X using the color camera mounted on a Leica DM6000 Widefield microscope.

### Immunofluorescence Assay on intestinal cryosections

Sections were thawed and incubated in PBS Triton 0.1% BSA 1% (hydration buffer), followed by blocking in PBS Triton 0.1% BSA 10% for 45 mins each. Antibody staining was carried out in hydration buffer. Primary antibodies were added for 2 hours, and for this study included rat anti- HA (1:500, Roche clone 3F10, #11867423001), LDH (1:200, kind gift from the Zhu Lab (Jilin University, China and Texas A&M University, USA) and rabbit anti α-tubulin (1:500, ABCD antibodies, #ABCD_AA345). Sections were washed three times (5 min incubation each time) in hydration buffer and incubated with secondary Alexa Fluor antibodies for 1h30 (1:500, Alexa Fluor 488, 568 or 647, Thermo Fisher Scientific). Sections were then incubated in hydration buffer containing Hoechst for 30 minutes (1:10000, Thermo Fisher Scientific #H3570), and then washed three times in hydration buffer (5 min each). Slides were mounted with Fluorogel (Electron Microscopy Science, Hatfield, PA).

### U-ExM on intestinal cryosections

Histology was performed as described above, then sections were rehydrated with hydration solution for 45 min and monomer solution (18% sodium acrylate, 10% acrylamide, 0.1% *N,N*ʹ- methylenebisacrylamide in PBS) was added to the slides for 90 min. Gelation was initiated by adding to the sections a polymerization mix composed of 90% monomer solution, 5% tetramethylethylenediamine (TEMED, 10% stock), and 5% ammonium persulfate (APS, 10% stock), in a total volume of 40 µl. Gels were polymerized on ice for 5 min, followed by overnight incubation at 37°C in a humidified chamber. To release the gels, a razor blade was used to scrape off the gel from the slides. Gels were then released from the coverslips and denatured at 95°C for 90 min in denaturation buffer (200 mM SDS, 200 mM NaCl, 50 mM Tris, pH 9). The rest of expansion was done as described above. Primary antibodies include rat anti-HA (1:500, Roche clone 3F10, #11867423001), rabbit anti α-tubulin (1:500, ABCD antibodies, #ABCD_AA345). Secondary antibodies include anti-rat 488 (1:500, Thermo Fisher Scientific # A-11006), Atto 647N NHS ester (stock in DMSO at 2 mg/mL, 1:200, Sigma-Aldrich #18373) and Hoechst (1:1000, Thermo Fisher Scientific #H3570).

### Fluorescence microscopy and image processing

Fluorescence images were acquired using a Leica DM6000B widefield microscope (coverslips) or a Leica STELLARIS confocal microscope (coverslips, tissue sections, and U-ExM gels). Image deconvolution was performed using Leica LAS X software. In HCT8 cells, parasite quantification, male gamete formation and egress were assessed using Hoechst, HA, and VVL staining. In infected mouse tissues, attachment and zygote quantification were assessed using Hoechst, male EGFP (Histone-H2B-EGFP driven by the male specific promoter cgd4_4380) and LDH staining. Male gamete nuclei dimensions, GGC ring sizes and the nuclear cap dimensions were measured using the “Measure Points” tool in Imaris 9.9 software. All additional image processing and visualization were conducted in ImageJ.

### Protein structure prediction and analysis

Protein sequences were aligned using MAFFT. GGC tertiary protein structures were predicted using the AlphaFold server (AlphaFold 3 model)^113^. Signal peptides were removed from protein sequences prior to prediction. Structural alignment was performed using the MatchMaker tool in ChimeraX 1.6.1. For structural similarity search, AlphaFold-predicted PDB files were uploaded to the Foldseek Search Server^114^ (version 10-941cd33). Eight databases (AlphaFold/Proteome, AlphaFold/Swiss-Prot, AlphaFold/UniProt50, BFMD, CATH50, GMGCL, MGnify-ESM30, PDB100) were searched using default parameters (3Di/AA mode).

### Attenuated infection and challenge

For vaccination infections, mice were infected with 5,000 or 50,000 HAP2-DiCre oocysts by oral gavage. Mice were treated with 10mg/kg rapamycin via oral gavage on D0-4 of infection, with the first dose given 4-5 hours prior to infection. Naïve controls were administered rapamycin only. Rapamycin was kept at a 50 mg/mL stock in ethanol and diluted in milliQ water immediately prior to treatment. 30 days after the mice had cleared the vaccine infection, mice were rechallenged with the 50k of wild type *C. parvum* or mouse-adapted *C. parvum.* To quantify fecal oocyst shedding, 20 mg of pooled cage feces was suspended in 1mL lysis buffer (deionized H_2_O with 50mM TrisHCl, 2mM DTT, 2 mM EDTA, 10% glycerol, 1% Triton). Samples were shaken with glass beads for 5 min, then combined in a 1:1 ratio with Nano-Glo® Luciferase solution (Promega, #N1150). A Promega GloMax plate reader was used to measure luminescence. Pooled samples were used because previous studies have demonstrated that mice within each cage are equally infected^115^.

### Antibody Treatments

For CD4 depletion, mice were treated with 400 µg of anti-CD4 (Assaygenie, clone: GK1.5, #IVMB0060) or isotype control (BioXCell, clone: LTF-2, #BE0090) via intraperitoneal injection three days before and on the day of the rechallenge infection.

### Quantification and statistical analysis

Statistical analyses were performed using R (version 4.5.2) and GraphPad Prism (version 11.0.2). Data are presented as mean ± standard deviation (SD). The statistical test used is indicated in figure legends.

---

**Figure S1:**
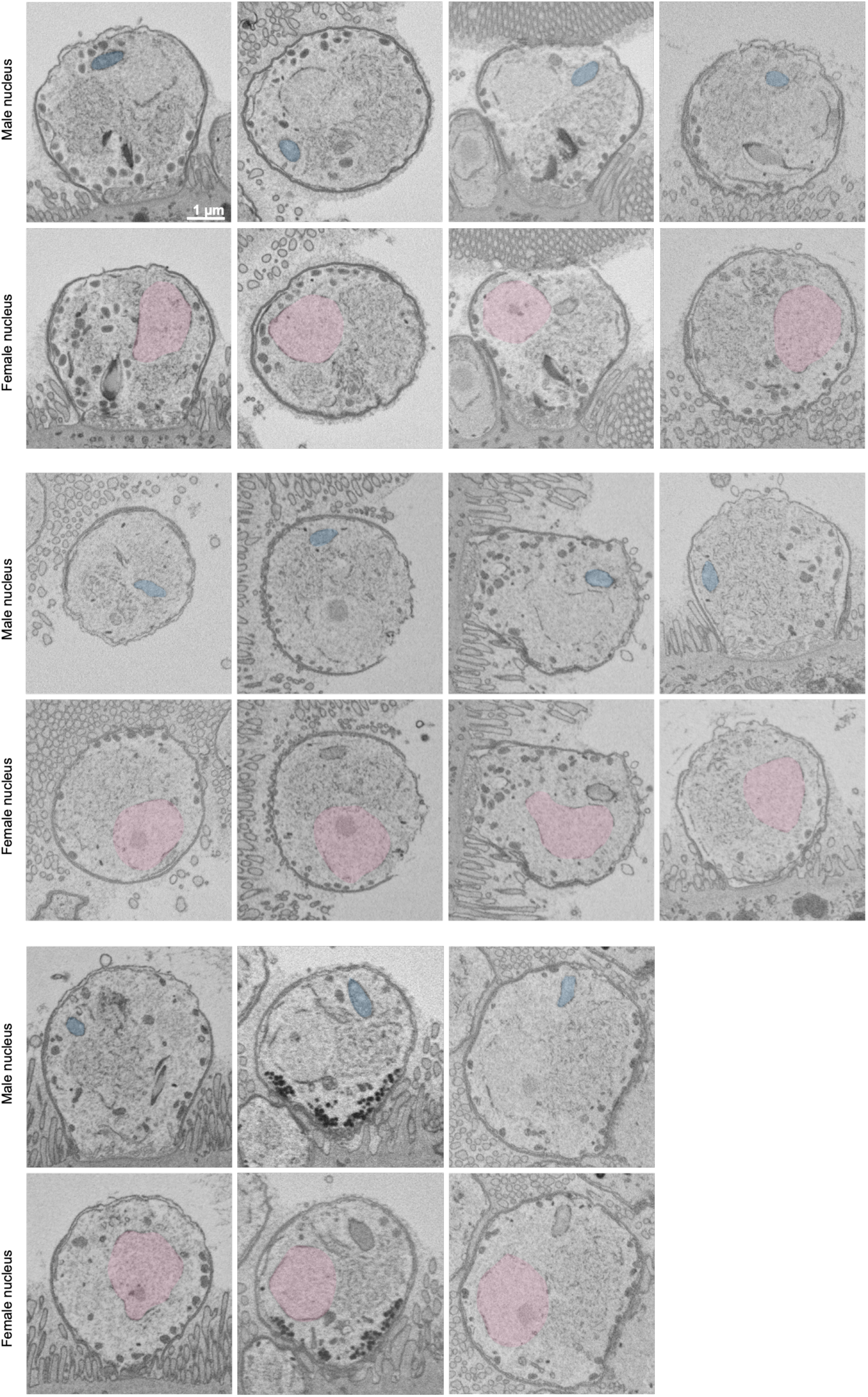
Eleven zygotes were identified by serial block-face scanning electron microscopy. Serial block-face scanning electron microscopy of intestinal sections from *C. parvum* infected mice showing the complete set of 11 zygotes found in this study. Male nuclei are highlighted in blue, female nuclei in pink. Scale bars = 1 µm.

**Figure S2:**
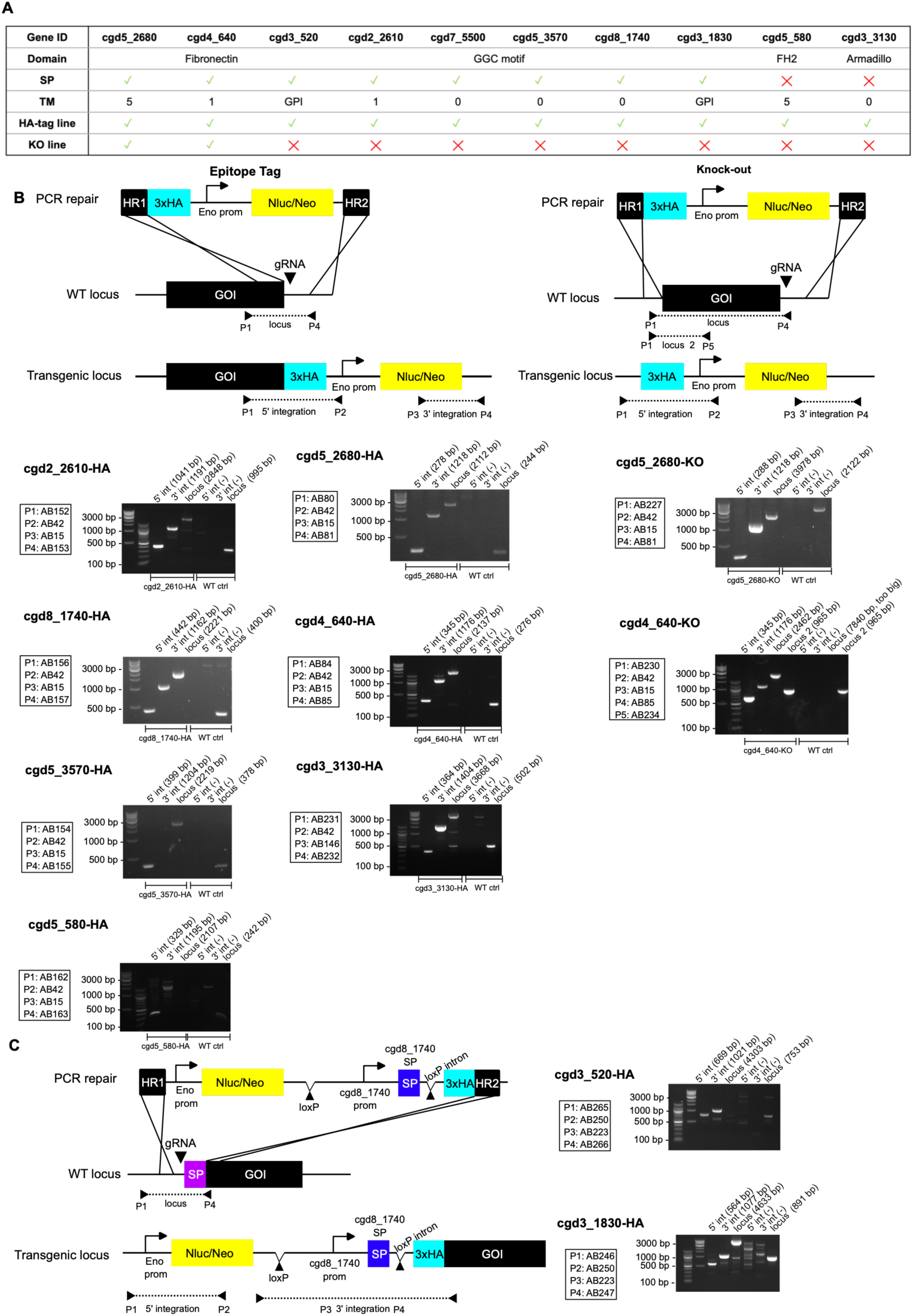
Generation of HA-tagged and knock-out strains of our 10 prioritized candidates. (**A**) Description of our 10 candidate proteins. SP: signal peptide; TM: transmembrane domain; GPI: glycosylphosphatidylinositol anchor. Whether a HA-tagged or KO line could be generated is specified by a ✓ (generated) or a ╳ (not generated). (**B**). Schematic maps of gene targeting resulting in HA-epitope-tagged (left) or deleted genes (right), with PCR analysis. (**C**). Generation of N-ter tagged strains, with PCR analysis. The promoter and signal peptide of cgd8_1740 were used and are in between loxP sites. The loxP after the signal peptide is in an intron. HR: homology region, Nluc: Nanoluciferase, Neo: neomycin resistance (Neo), GOI: gene of interest.

**Figure S3:**
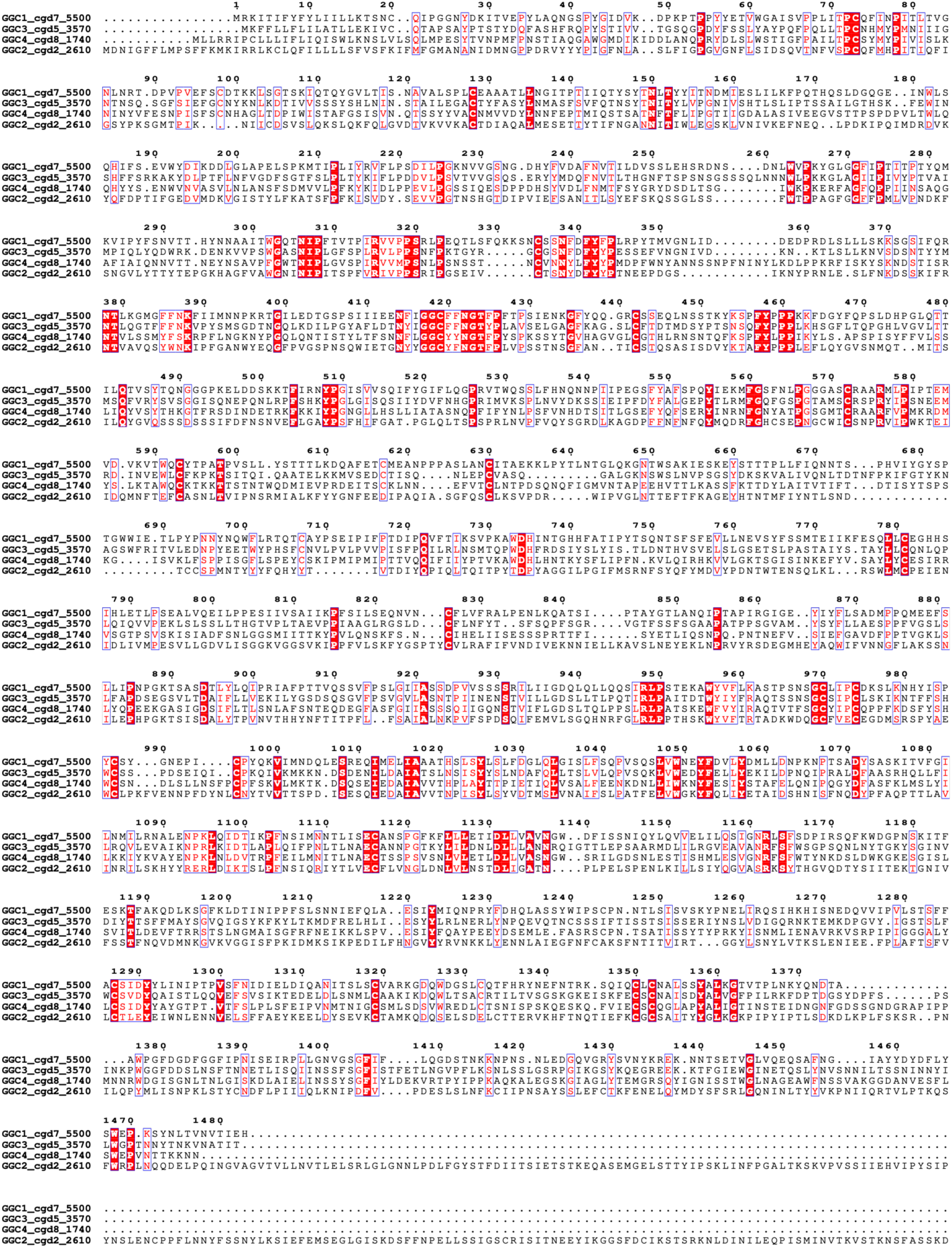
Multiple sequence alignment of GGC1-4 shows low sequence similarity. Note the motif GGC%%NGT%P (position 417) which gives this family of protein its name. If this motif possesses a role is unknown.

**Figure S4:**
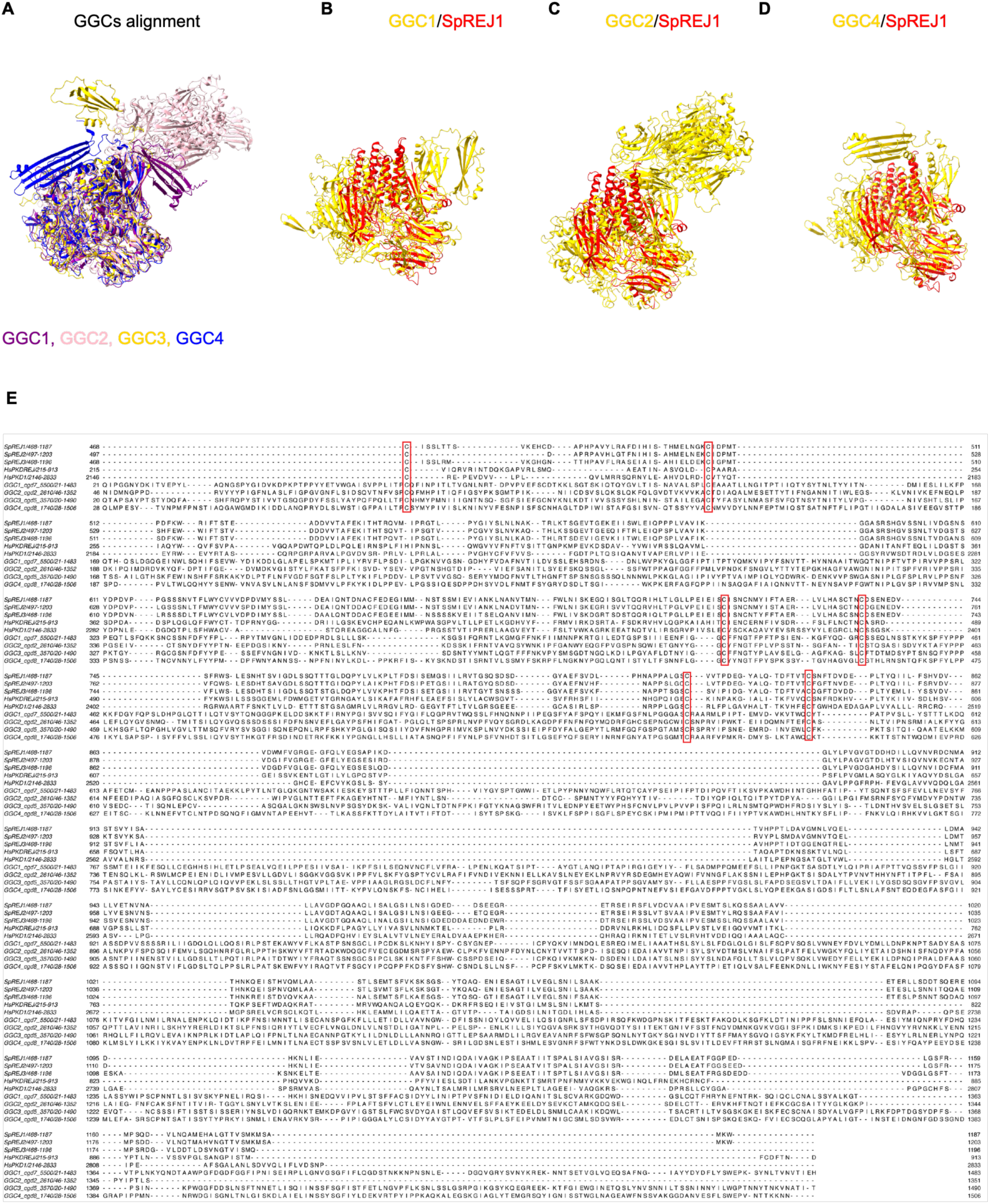
GGC proteins show structure and sequences similarities with REJ-domain containing proteins. (**A**) Overlay of AlphaFold structural predictions for GGC1, GGC2, GGC3 and GGC4 (omitting the signal peptide). RMSD: GGC1/GGC2: 1.336 Å across 480 residues; GGC1/GGC3: 1.081 Å across 893 residues; GGC1/GGC4: 1.183 Å across 744 residues; GGC2/GGC3: 1.270 Å across 473 residues, GGC2/GGC4: 1.262 Å across 523 residues; GGC3/GGC4 1.166 Å across 743 residues. (**B-D**). Structure alignment of the REJ domain of SpREJ1 (SpREJ1_468-1187_) with GGC1 (RMSD: 1.134 Å across 55 residues) (**B**), GGC2 (RMSD: 1.168 Å across 36 residues) (**C**) and GGC4 (RMSD: 1.417 Å across 53 residues) (**D**). (**E**). Multiple sequence alignment of GGC1-4 with the REJ domains of SpREJ1-3, HsPKDREJ and HsPKD1 reveals the presence of the conserved 6-cysteines, typical of REJ domains, in GGC proteins. The signal peptide of GGC1-4 were removed and only the first unit of GGC2 was aligned (determined by visualization of the AlphaFold prediction, amino acid 46-1352).

**Figure S5:**
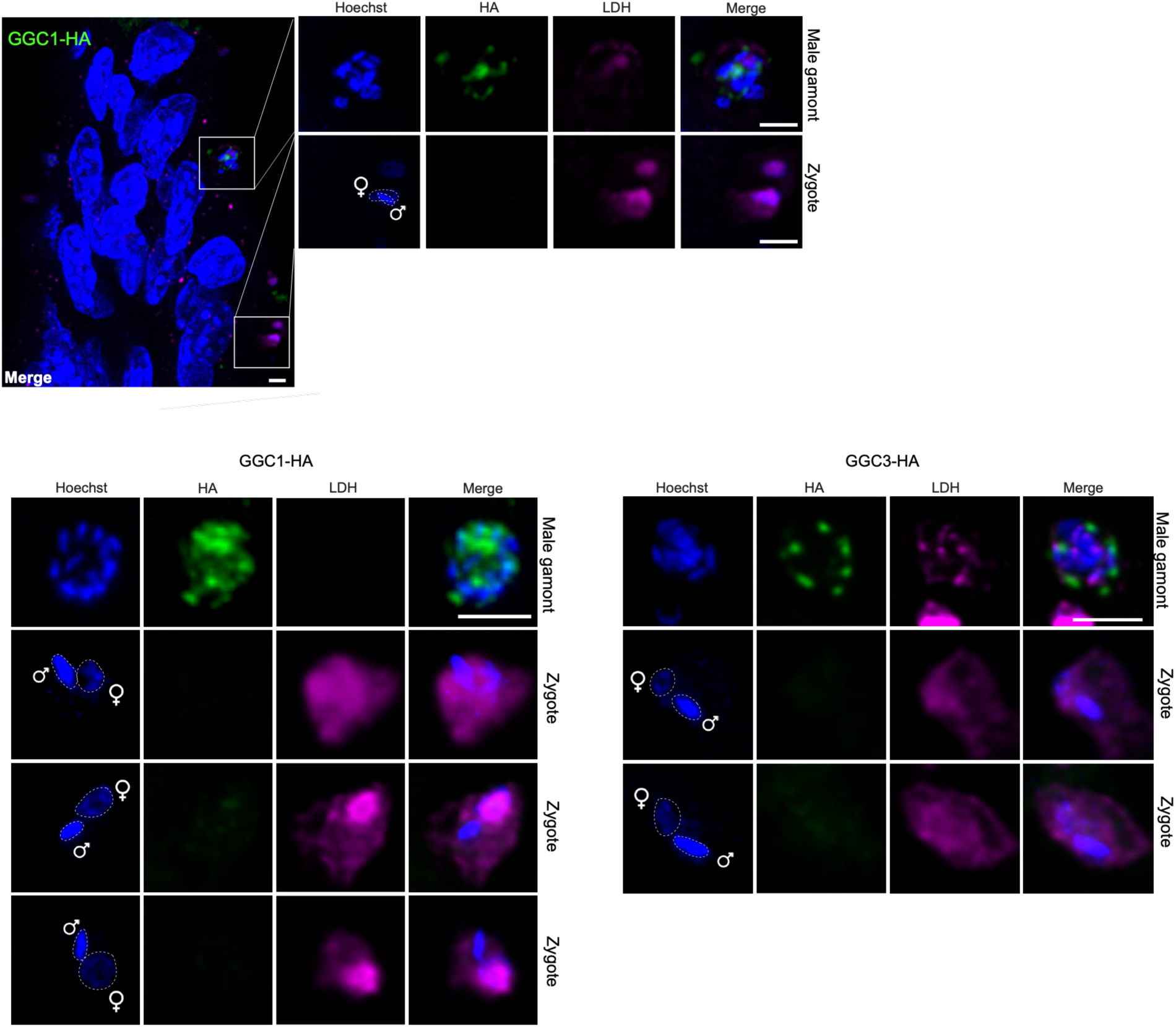
GGC proteins are present in male gamonts but absent from zygotes. Representative intestinal sections from GGC1-HA and GGC3-HA infected mice, with inset showing a male gamont and a zygote. The male and female nuclei of the zygotes are highlighted. Magenta: lactate dehydrogenase (LDH) antibody used as pan-parasite stain. Scale bars: 2 µm.

**Figure S6:**
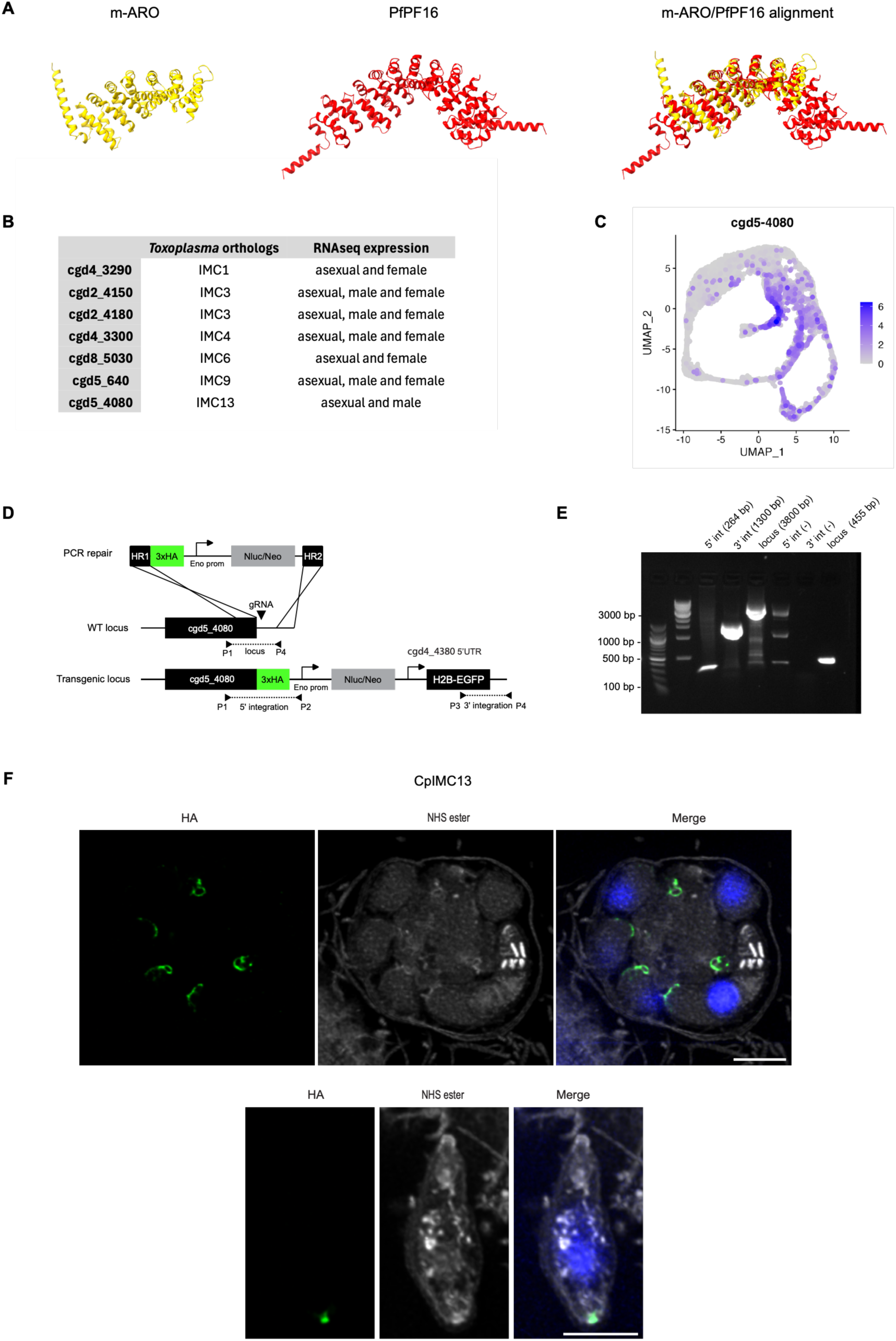
IMC13 is conserved in *Cryptosporidium.* (A) Left: AlphaFold prediction of m-ARO and PF16 from *Plasmodium.* Right: Structure alignment of m-ARO and PfPF16. See Video S4 for movie of the alignment. RMSD: 1.063 Å across 79 residues. (B) Conservation of *Toxoplasma* alveolins in *Cryptosporidium* and their expression in the *C. parvum* single cell transcriptome^20^. (**C**) UMAP representation of CpIMC13 (cgd5_4080) in the *C. parvum* single cell transcriptome. (**D-E**). Generation of a C-ter tagged strain of CpIMC13 with the schematic map in (**D**) and PCR analysis in (**E**). (**F**). U-ExM images of CpIMC13-HA in developing meronts (top) and egressed merozoites (bottom). Note the presence of the protein in the basal complex. Hoechst (blue) and N-hydroxysuccinimide ester (NHS) protein counterstain in white. Scale bar: 5 µm. Expansion factor: 4.25.

**Figure S7:**
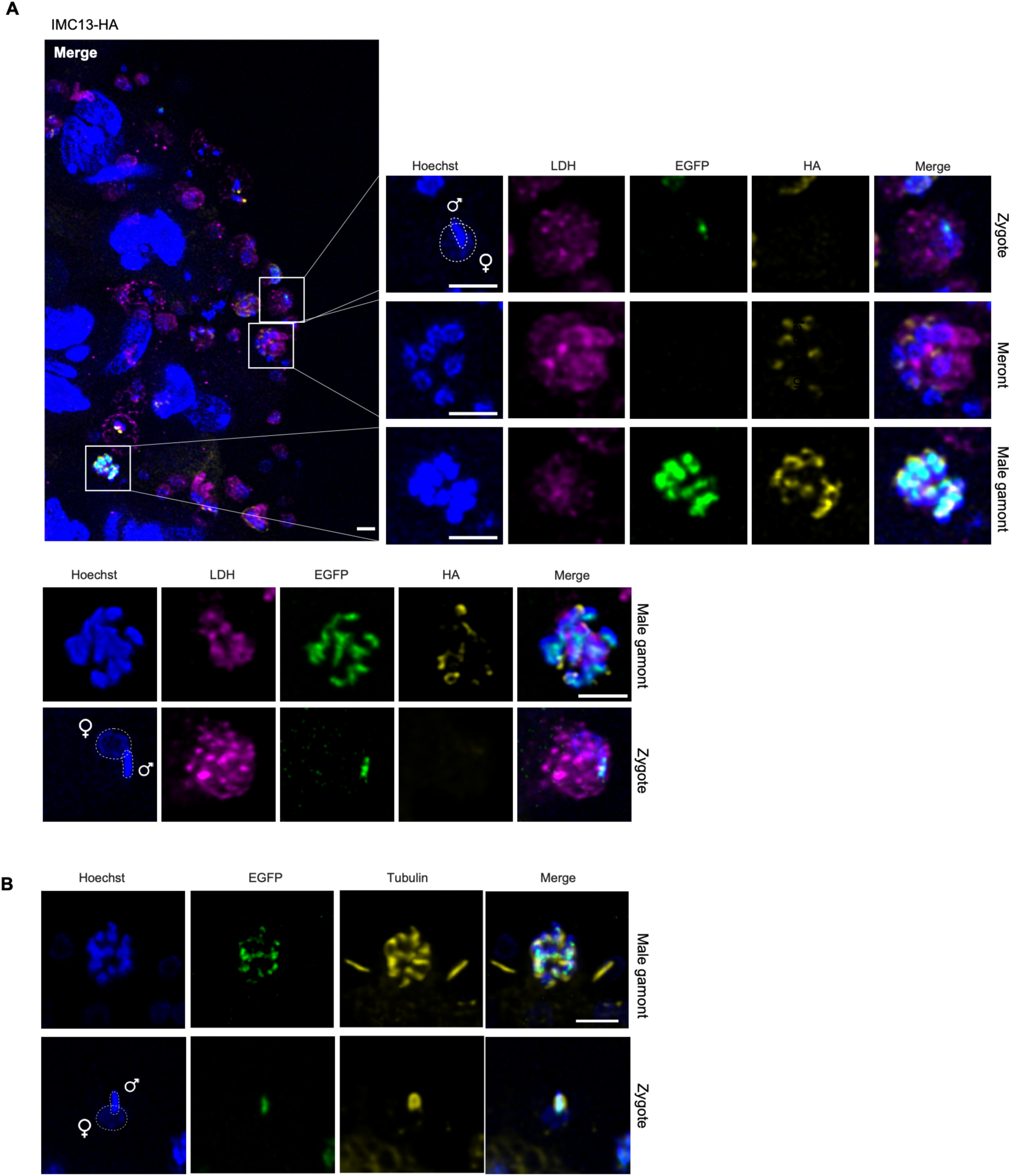
Microtubules, but not CpIMC13, are internalized in the zygote during fertilization. (A) Representative intestinal sections from IMC13-HA infected mice, with inset showing a zygote, a male gamont and a meront. The male and female nuclei of the zygote are highlighted. Green: EGFP, male nuclei, Magenta: LDH, lactate dehydrogenase antibody used as pan-parasite stain. Scale bars: 2 µm. (**B**) Representative intestinal sections from infected mice showing a male gamont and a zygote stained with an α -tubulin antibody. A transgenic *C. parvum* strain expressing male EGFP in the nuclei was used. Scale bar: 2 µm.

**Figure S8:**
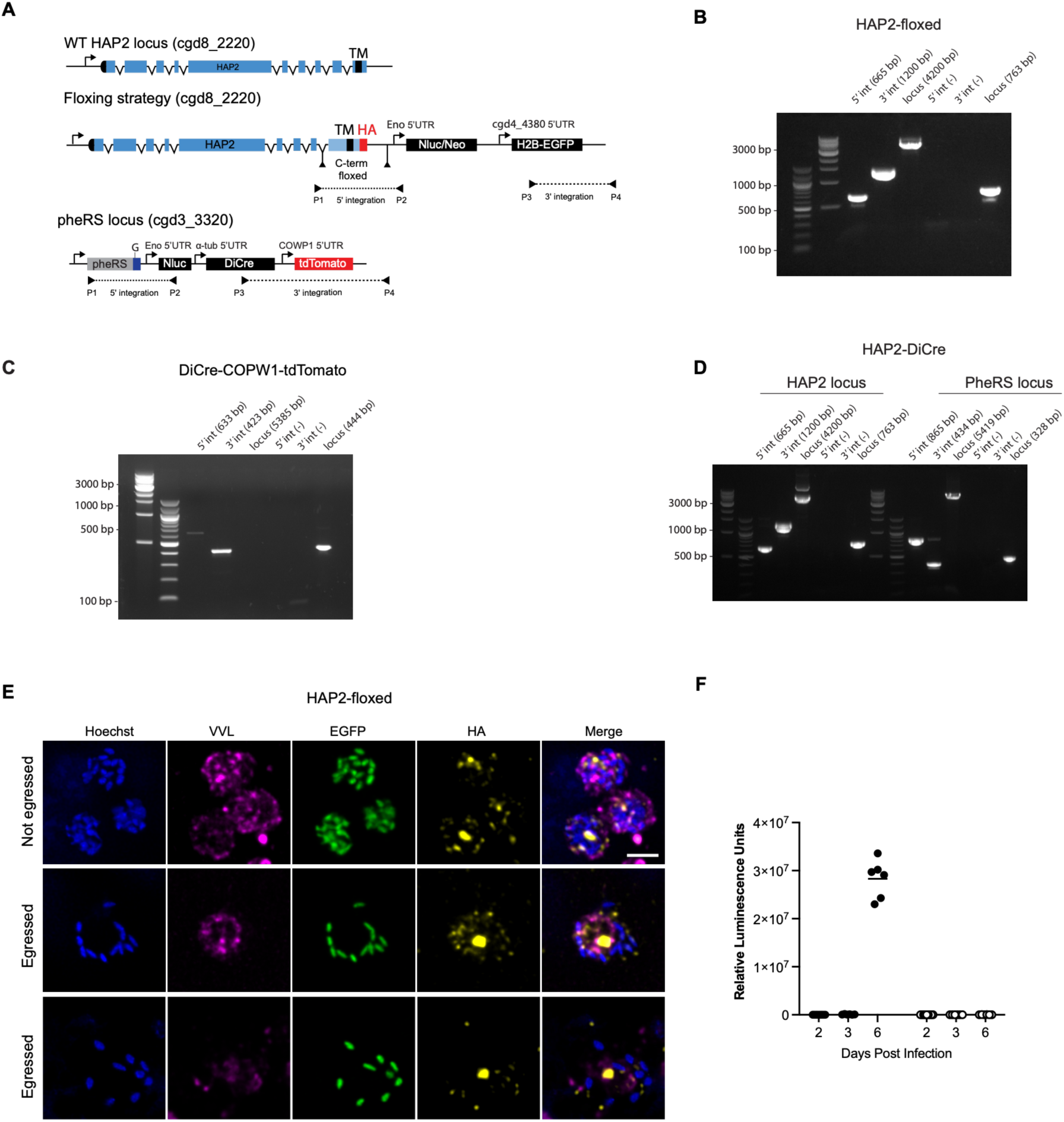
HAP2-DiCre strain generation and supplementary information related to. Figure 6. (A) Top: Genetic strategy to replace the transmembrane domain of HAP2 with a recodonized sequence, including an HA epitope tag and flanking loxP sites (black triangles). The internal loxP site was placed within an intron. Nanoluciferase (Nluc), neomycin resistance (Neo) and a male-specific expressing Histone2B-EGFP cassette were inserted downstream of the gene. Bottom: The pheRS locus was targeted to introduce the DiCre recombinase, the Nluc cassette, and a female-specific expressing tdTomato. (**B-C**). PCR analysis showing proper integration of the floxed construct in the HAP2 locus (**B**) and of the DiCre cassette in the pheRS locus (**C**). Primers annealing location are shown in A. The resulting HAP2-floxed and DiCre transgenic strains were crossed, and dual drug selection was used to isolate HAP2-DiCre. PCR analysis showing proper integration of the two construct is shown in (**D**). (**E**). Representative images of HAP2-floxed infected HCT8 cells showing male gamonts before egress (top) and egressed male gametes (bottom). VVL: antibody used as pan-parasite stain, EGFP: male nuclei. (**F**). Luminescence assay of intestinal punch biopsies from infected mice treated with vehicle or rapamycin. Each dot represents one biopsy, 12 mice total.

**Video S1: SBF-SEM volumes of the female gamete development, related to Figure 1B**. Serial block-face scanning electron microscopy of intestinal sections from *C. parvum* infected mice. Animation through 4 SBF-SEM Z-stacks: a female gamete type I, type II, an oocyst with a single nucleus and an oocyst with 4 nuclei. Nuclei of females are highlighted in pink; nuclei of oocysts are highlighted in purple.

**Video S2: SBF-SEM volumes of male gamonts, related to Figure 1A**. Serial block-face scanning electron microscopy of intestinal sections from *C. parvum* infected mice. Animation through 2 SBF-SEM Z-stacks of two mature male gamonts.

**Video S3: SBF-SEM volumes of attached male and intracellular zygotes, related to Figure 1D-E**. Serial block-face scanning electron microscopy of intestinal sections from *C. parvum* infected mice. Animation through a male gamete attached to a host cell surrounding a female gamete and 3 SBF-SEM Z-stacks of zygotes. Nuclei of females are highlighted in pink; nuclei of males are highlighted in blue.

**Video S4: Alignment of AlphaFold predicted structure related to Figure 4A-B, Figure S4A, Figure S6A.** Animation through 3 protein structure alignment: 1) Structure alignment of GGC1-4; 2)

Structure alignment of GCC3 with the REJ domain of SpREJ1 ; 3)) Structure alignment of m-ARO with PfPF16 from *Plasmodium.* Alignments were done using the Matchmaker tool of Chimera.

## Acknowledgements

This work was supported in part by grants from the National Institutes of Health R37-AI112427 and R01AI127798 to BS; R01AI148249 and U01AI163671 to CAH and BS and F30AI186259 to MM; EMBO Fellowship ALTF 1145-2021 to ACB, a Mark and Pamela Winter Infectious Disease Fellowship to AMD, and support from the Robert Koch Institute to TB, ML, and CK. The Leica STELLARIS was purchased using NIH funding grant S10 OD032305-01A1. We thank Dr. Gordon Ruthel at the PennVet Imaging Core (RRID: SCR_022764) for assistance. We thank J. Byerly and G. Buenconsejo for support in animal experimentation, and A. Cohen, C. Nötzel, A. Guerin and S. Shaw for transgenic strains, ongoing discussion and support.

